# A Study of Calibration as a Measurement of Trustworthiness of Large Language Models in Biomedical Research

**DOI:** 10.1101/2025.02.11.637373

**Authors:** Rodrigo de Oliveira, Matthew Garber, James M Gwinnutt, Emaan Rashidi, Jwu-Hsuan (Shantina) Hwang, William Gilmour, Jay Nanavati, Khaldoun Zine El Abidine, Christina DeFilippo Mack

## Abstract

**Objectives:** To assess the calibration of 9 large language models (LLMs) within biomedical natural language processing (BioNLP) tasks, furthering understanding of trustworthiness and reliability in real-world settings.

**Materials and Methods:** For each LLM, we collected responses and corresponding confidence scores for all 13 datasets (grouped into 6 tasks) of the Biomedical Language Understanding & Reasoning Benchmark (BLURB). Confidence scores were assigned using 3 strategies: Verbal, Self-consistency, Hybrid. For evaluation, we introduced Flex-ECE (Flexible Expected Calibration Error): a novel adaptation of ECE that accounts for partial correctness in model responses, allowing for a more realistic assessment of calibration in language-based settings. Two post-hoc calibration techniques—isotonic regression and histogram binning—were evaluated.

**Results:** Across tasks, mean calibration ranged from 23.9% (Population-Intervention-Comparison-Outcome extraction) to 46.6% (Relation Extraction). Across LLMs, Medicine-Llama3-8B had the best mean overall calibration (29.8%); Flan-T5-XXL had the highest ranking on 5/13 datasets. Across strategies, self-consistency (mean: 27.3%) had better calibration than Verbal (mean: 42.0%) and Hybrid (mean: 44.2%). Post-hoc methods substantially improved calibration, with best mean calibrated Flex-ECEs ranging from 0.1% to 4.1%.

**Discussion:** The poor out-of-the-box calibration of LLMs poses a risk to trustworthy deployment of such models in real-world BioNLP applications. Calibration can be improved post-hoc and is a recommended practice. Non-binary metrics for LLM evaluation such as Flex-ECE provide a more realistic assessment of trustworthiness of LLMs, and indeed any model that can be partially right/wrong.

**Conclusion:** This study shows that out-of-the-box calibration of LLMs is very poor, but traditional post-hoc calibration techniques are useful to calibrate LLMs.

## LAY SUMMARY

Large-language models (LLMs) are increasingly being used within biomedical research, yet quantitative benchmarking of the trustworthiness and reliability of LLMs is often not performed. This is a risk, especially in healthcare when the output of these models is used to guide decision making.

Confidence scores are metrics that can assess the trustworthiness, or reliability, of an LLM’s output. For calibration scores to be reliable, the model must be well calibrated, meaning that the model’s confidence and accuracy are broadly aligned. A poorly calibrated model may provide inaccurate responses with high confidence, leading to downstream errors and potentially significant risk to healthcare users.

This study assessed the calibration of 9 LLMs across 13 datasets critical for biomedical research. There was substantial variation in calibration across LLMs; however, even the best performing LLMs were approximately 30% off target, indicating the LLMs are poorly calibrated for biomedical tasks. Luckily, traditional and inexpensive correction methods were demonstrated to improve the LLMs’ calibration.

This study demonstrates a tailored strategy matching LLMs with biomedical tasks and applying appropriate corrections is required to achieve optimal calibration, and essential for decision-making in a healthcare setting; a “one-size fits all” approach is not sufficient.

## BACKGROUND AND SIGNIFICANCE

The increasing deployment of large language models (LLMs) in high-stakes domains such as healthcare, where precise and reliable information is critical, has prompted growing scrutiny over their trustworthiness. Biomedical Natural Language Processing (BioNLP) applications, which encompass tasks such as disease identification, medical decision support, and information extraction, require human oversight for interpretation. Confidence metrics can enhance this oversight by serving as proxies for trustworthiness, by associating a response with a probability of the validity of the response.

However, confidence metrics are only reliable if the model is well calibrated. Calibration is assessed by measuring the degree of alignment between the model’s predicted probabilities and true probabilities (e.g. accuracy) over a test set [1]. For example, if a model predicts a 75% chance of cancer in radiographs, we expect 75% of those images to indicate cancer. Calibration ultimately informs developers and users of a model whether predicted probabilities can be interpreted as *confidence* scores.

Model probabilities as confidence scores are useful for downstream decisions, and are necessary in high-stakes environments, such as in healthcare. For instance, if a model is well-calibrated, we might trust its predictions without human intervention when confidence scores are extreme (e.g. >=95% or <=5%), while opting to manually verify predictions with confidence scores outside of the ranges. When a model is *miscalibrated*, we cannot trust predicted probabilities and would need to manually review all predictions. Measuring calibration reveals whether the LLM’s confidence in its responses is accurate, indicating whether it is overconfident, under-confident, or *just right*, as illustrated in Figure 1.

**Figure 1:**
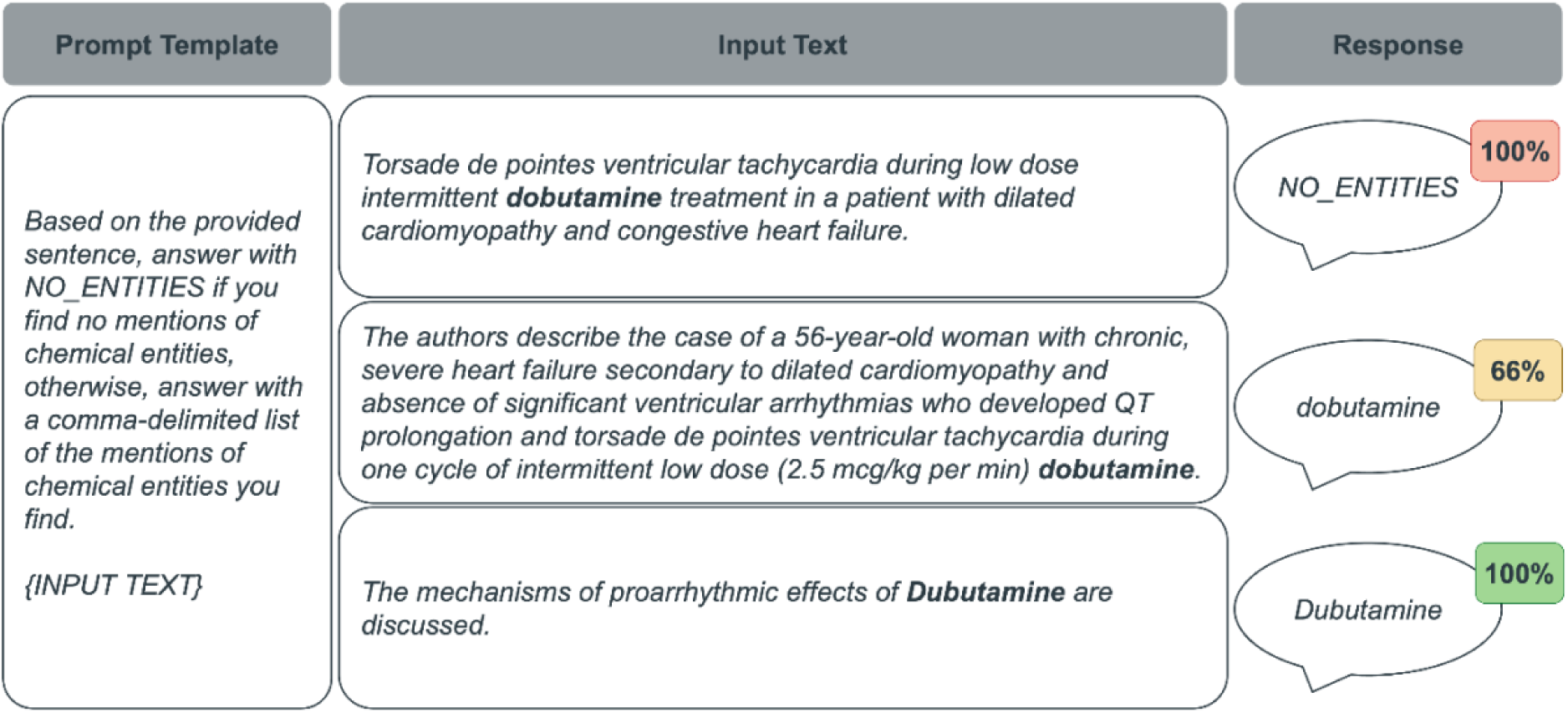
Examples of GPT-3.5 being over-confident (top), under-confident (middle) and‘just right’ (bottom) when identifying chemicals (BC5-chemical). Highlighted snippets are the chemicals to identify in each text, and percentages are the confidence scores associated with each response, which in this case were assigned using the self-consistency strategy (more on strategies below).

Despite recent advancements, LLMs frequently struggle with overconfidence in their predictions [2], especially in BioNLP. This is largely due to a lack of domain-specific training, as the most popular LLMs are trained on general-domain data sources like social media and Wikipedia [3]. There is a growing body of research focused on calibration in LLMs, yet there remains a significant gap in applying these findings to BioNLP tasks for healthcare research [4,5]. This paper aims to address this gap by first assessing the degree of calibration of various LLMs in various BioNLP tasks. Having quantified the degree of calibration—or miscalibration—of LLMs in BioNLP, we then proceed to evaluate the effectiveness of different techniques to mitigate miscalibration.

## OBJECTIVES

The objectives of the study are to quantify:

1. The degree to which LLMs are calibrated for BioNLP tasks.
2. The utility of traditional calibration techniques to mitigate miscalibration of LLMs in the BioNLP domain.

Specifically, we wish to address the following research questions (RQs):

**RQ1** How calibrated are state-of-the-art LLMs in the BioNLP domain?

**RQ1.1** Is there a relationship between calibration and *family* of LLMs?

**RQ1.2** Is there a relationship between calibration and *family* of BioNLP tasks?

**RQ1.3** Is there a relationship between calibration and confidence assignment strategy?

**RQ2** Can post-hoc calibration techniques mitigate LLM miscalibration?

## MATERIALS AND METHODS

Figure 2 illustrates how we systematically assessed the calibration of a diverse set of LLMs. Below sections delve into details of our approach.

**Figure 2:**
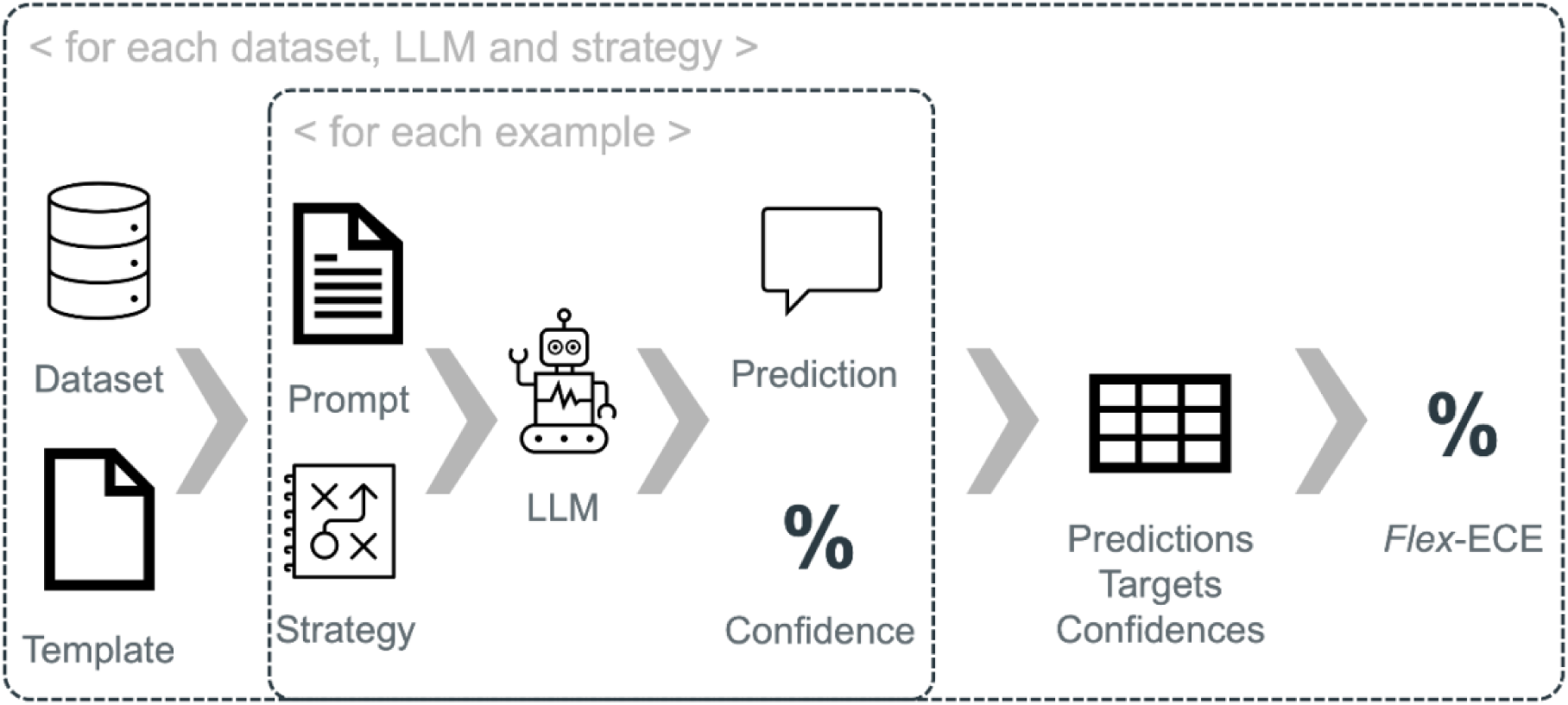
A Flex-ECE score is computed for an LLM with respect to a particular dataset (task) and confidence assignment strategy.

### BLURB Benchmarking Datasets and Healthcare-Based Tasks

The Biomedical Language Understanding & Reasoning Benchmark (BLURB)^1^ is a comprehensive benchmark for BioNLP with thirteen publicly available datasets, grouped in six tasks. BLURB has been thoroughly described previously [6], but Figure 3 provides an illustration of each task for ease of reading.

**Figure 3:**
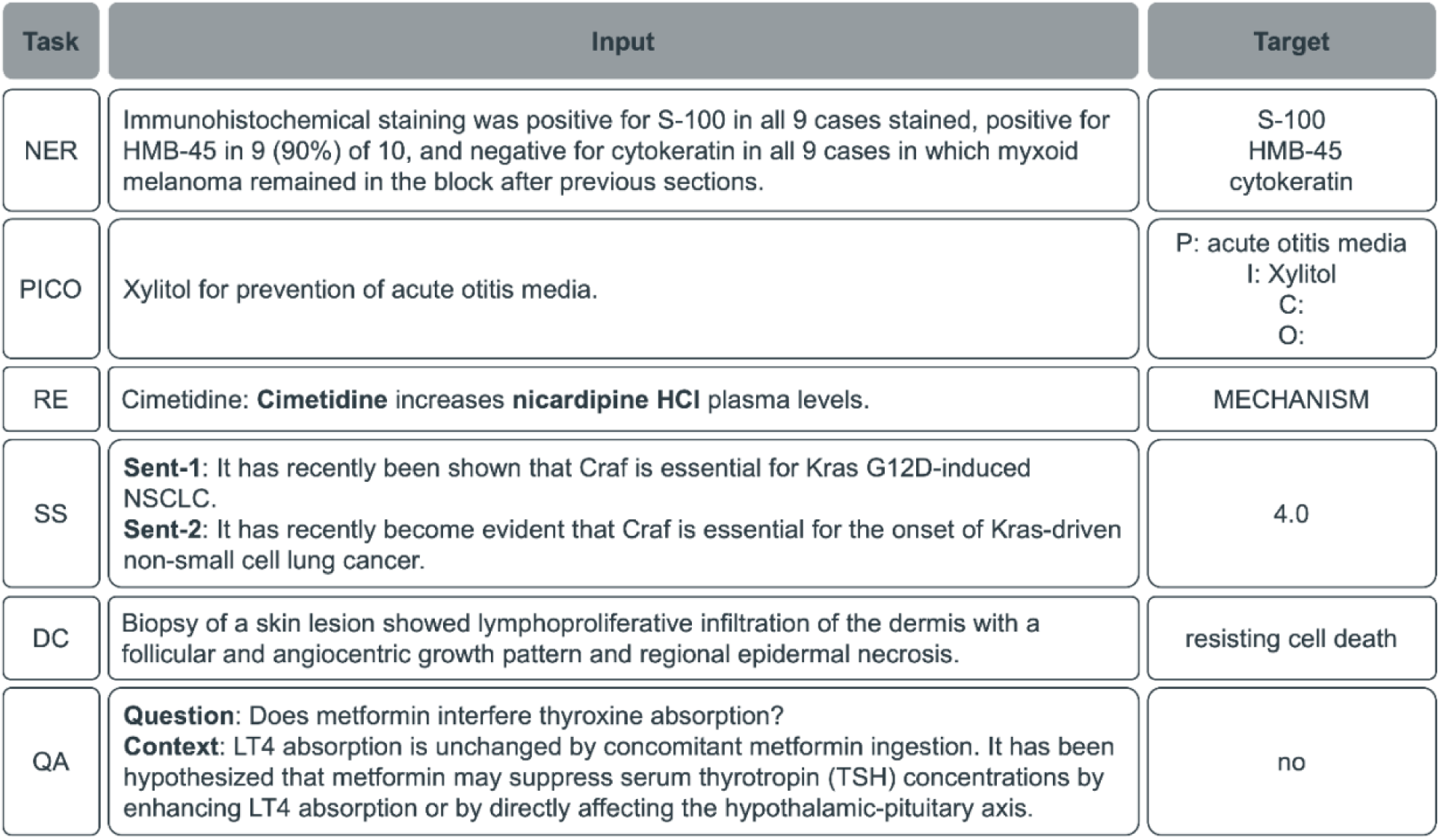
BLURB datasets contain an input text, which we inject in prompt templates, and a target (correct) label, which is used to compute similarity (prediction vs. target), and ultimately calibration (confidence vs. similarity).

### Models & Prompting

Our study included a diverse set of nine LLMs, differing in both size of learnable parameters (stratified into small [≤10 billion parameters] and large [>10 billion parameters]) and domain specificity of the data used to train (or fine-tune) models, resulting in 4 “families” of models:

- General domain:

`o`**Small:** Zephyr-7B-Beta [7], Llama-3-8B-Instruct [8]
`o`**Large**: Flan-T5-XXL [9], Yi-1.5-34B-Chat [10], GPT-3.5-Turbo [11], GPT-4[11]
- Biomedical domain:

`o`**Small**: Meditron-7B [12], Medicine-Llama3-8B [13,14]
`o`**Large**: MedLLaMA-13B [15]

All above models are *generative* LLMs, which means we need to *prompt* them with natural language (text). We submit a unique prompt for each example of each dataset to LLMs by programmatically filling a *prompt template*^2^ with the input text of each example. The dataset-specific prompt templates contain instructions for what the LLM should do for all examples of the given dataset.

### Confidence Assignment Strategies

As aforementioned, we require (*potential*) confidence scores for each response, so that we can compute a model’s calibration. Assigning a confidence score to an LLM’s response is not as trivial as it is with traditional classifiers, whose responses *are* confidence scores. LLMs are effectively next word^3^ predictors, so the only probabilities we know for certain are the probabilities of each word in the response given the previous words, but not of the response as a whole, or of the claim in the response [16].

In line with [2], three confidence assignment techniques^4^ have been explored:

- **Verbal**: Consists of simply including a request in the prompt to return a confidence score in addition to the response; more sophisticated verbal strategies have not proven to yield better confidence scores [2,16].
- **Self-consistency**: Consists of sampling multiple responses for the same prompt^5^, then computing the degree of *similarity* between the several candidate responses and the original response. The final confidence score is the average of similarity scores.
- **Hybrid**: Consists of combining both above strategies, by eliciting multiple verbal confidence scores, then boosting or penalising the initial verbal score depending on the consistency of the responses.

### Evaluation

Traditional calibration metrics such as *Expected Calibration Error* (ECE) [1] assume that models’ predictions are entirely right or wrong, so that true probabilities become the ratio of right answers for a given test set, i.e. *accuracy*. Treating LLM responses as entirely right or wrong is not desirable, often not feasible [17].

This study thus proposes a *flexibilization* of ECE, dubbed ***Flexible ECE*** or ***Flex-ECE***^6^ for short, where model accuracy is the average *similarity* between generated and target responses for a given test set; the binning approach of Flex-ECE is the same as in conventional ECE (we use 5 bins for all experiments). A task-specific similarity function^7^ is employed, as illustrated in Figure 4.

**Figure 4:**
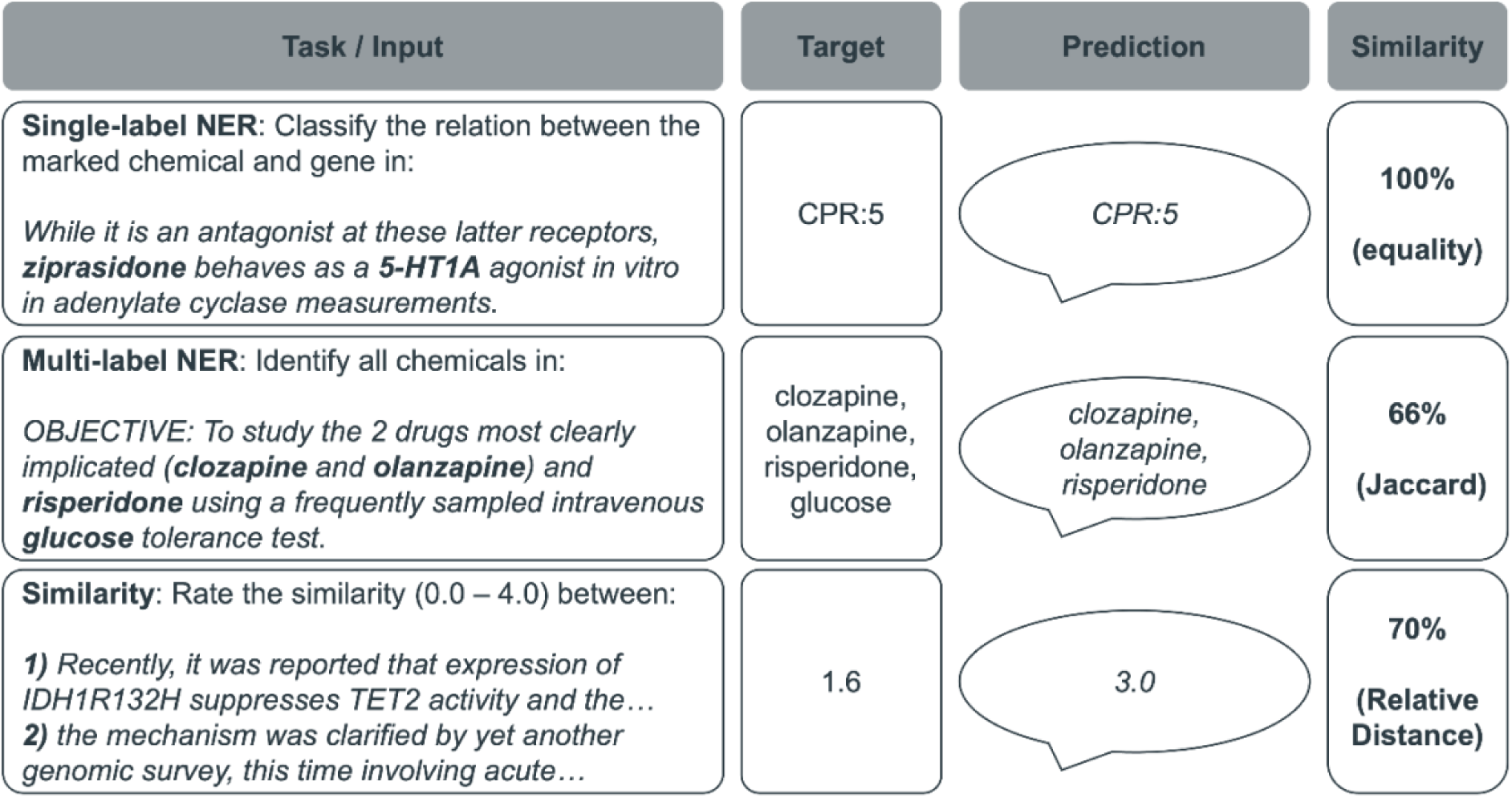
Task-specific similarity functions prevent us from severely punishing models in scenarios where being partially correct would be acceptable, by allowing correctness to be continuous (e.g. a percentage) rather than binary (i.e. right or wrong).

### Post-Hoc Calibration

In addition to quantifying the out-of-the-box degree of (mis)calibration of LLMs, we also investigated ways to mitigate miscalibration. We explore only post-hoc calibration techniques, i.e. we modify confidence scores only while leaving models intact, since re-training LLMs can be very costly [18]. We wish to first understand to what extent traditional, inexpensive calibration techniques can be used to extract more accurate confidence scores from LLMs. Two techniques were tested:

- **Isotonic regression**: used to learn the rescaling factor that arranges the observed set of probabilities in a stepwise, monotonically increasing line [19].
- **Histogram binning**: a stricter version of isotonic regression, where the steps (or bins) are pre-defined or are distributed so that each bin has an equal sample size [20].

To avoid overfitting, we fit each calibration method on a randomly selected subset of the train split of each BLURB dataset, using the uncalibrated confidences from the best performing confidence elicitation strategy for each model, as determined by mean Flex-ECE across datasets. As some calibration methods (particularly isotonic regression) are prone to overfitting when there is insufficient calibration data [21], we test the effect of varying the number of calibration samples (10, 50, 100, 500, and 1000).

### Infrastructure

We ran experiments one LLM at a time and utilized the following hardware to serve each self-hosted LLM (which excludes both GPT models): 8-32 x vCPUs, 32-64GB of RAM and 1 Nvidia GPU with 32-40GB of RAM for most LLMs, except for the two largest models (Yi-1.5-34B-Chat and Flan-T5-XXL), where we used 2 such GPUs. To execute experiment scripts, 2 x vCPUs with 8GB of RAM and no GPUs were used. Scripts were all written in Python 3.9, including our self-developed calibration metric *Flex-ECE*.

## RESULTS

The number of examples per task ranged from 20 (Sentence similarity) to 5038 (Gene recognition) (Table 1). A verbal confidence score was extracted (defined as a “success”) for 100% of examples for the majority of dataset-LLM combinations. Exceptions where <50% success was observed were: Medicine-Llama3-8B (Biomedical question answering with context task:17.0% success); Meditron-7B and Zephyr-7B-Beta (all tasks other than PICO extraction: <6.8% success); MedLLaMA-13B (Chemical recognition: 16.4%; Hallmarks of Cancer: 22.0%). When the Self-consistency and Hybrid strategies were used, all LLMs had 100% success rates, except GPT-3.5 and GPT-4 (≥99.8% across all datasets).

**Table 1:**
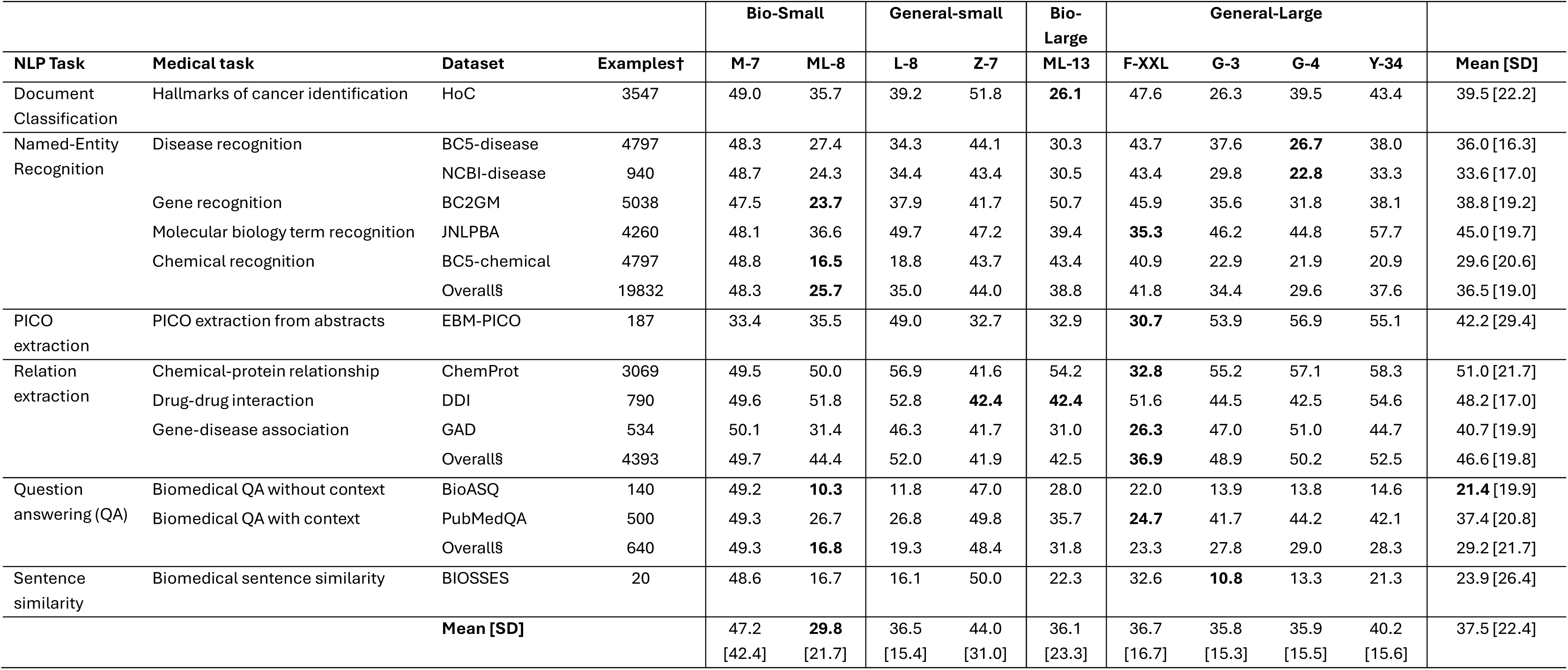
Mean Flex-ECE scores by dataset and model. The calibration scores in this table represent the mean of the Flex-ECE derived from each of the three confidence score elicitation strategies. Only task-model-strategy combinations with a success rate ≥50% are included. PICO: Population, Intervention, Comparator, Outcome; QA: Question asnwering; NLP: Natural language processing; SD: Standard deviation Models are abbreviated for ease of reading: F-XXL = Flan-T5-XXL; G-3 = GPT-3.5; G-4 = GPT-4; ML-13 = MedLLaMA-13B; ML-8 = Medicine-Llama3-8B; M-7 = Meditron-7B; L-8 = Meta-Llama-3-8B-Instruct; Y-34 = Yi-1.5-34B-Chat; Z-7 = Zephyr-7B-Beta. Bio-small: M7, ML-8; General-Small: Z-7, L-8; Bio-Large: ML-13; General-Large: F-XXL, Y-34, G-3, G-4 † Indicates the number of examples within each of the datasets (e.g. within the HoC dataset, there were 3547 different examples texts for LLMs to label with hallmarks of cancer) § The Overall rows indicate the mean of the calibration scores from each dataset for a given NLP task. For example, the Overall row of the Named-Entity Recognition section is the mean of the calibration scores from the BC5-disease, NCBI-disease, BC2GM, JNLPBA, and BC5-chemical datasets.

### RQ1: How calibrated are state-of-the-art LLMs in the BioNLP domain?

**Error! Reference source not found.**The calibration of LLMs was generally poor for tasks in the biomedical domain (Table 1). The Biomedical question answering without context task had the best calibration (mean: 21.4%), followed by the Sentence similarity task (mean: 23.9%). The Chemical-protein relationship extraction task (mean: 51.0%) had the worst calibration, followed by the Drug-drug interaction extraction task (mean: 48.2%).

Medicine-Llama3-8B had the best calibration across the datasets (mean: 29.8%; Table 1). Five of the LLMs had similar mean calibration scores across the datasets (means ranging from 35.8% to 36.7%; ordered by increasing score: GPT-3.5, GPT-4, MedLLaMA-13B, Meta-Llama-3-8B-Instruct, and Flan-T5-XXL). The LLMs with the worst calibration were Yi-1.5-34B-Chat (mean: 40.2%), Zephyr-7B-Beta (mean: 44.0%) and Meditron-7B (mean: 47.2%). Medicine-Llama3-8B was ranked as the best calibrated model for only 3 / 13 datasets whereas Flan-T5-XXL was ranked as the best calibrated model for 5 / 13 datasets.

### Document Classification

#### Hallmarks of cancer (HoC)

The LLMs with the best calibration were MedLLaMA-13B (mean: 26.1%) and GPT-3.5 (mean: 26.3%). The LLM with the worst calibration was Zephyr-7B-Beta (mean: 51.8%)

### Named-entity Recognition

#### Disease recognition (BC5-disease and NCBI-disease)

The LLMs with the best calibration were GPT-4 (mean = BC5-disease: 26.7%; NCBI-disease: 22.8%) and Medicine-Llama3-8B (mean = BC5-disease: 27.4%; NCBI-disease: 24.3%). Meditron-7B had the worst calibration (mean = BC5-disease: 48.3%; NCBI-disease: 48.7%).

#### Gene recognition (BC2GM

Medicine-Llama3-8B had the best calibration (mean: 23.7%). The mean score for GPT-4 was 31.8% and the scores for GPT-3.5, Meta-Llama-3-8B-Instruct and Yi-1.5-34B-Chat ranged from 35.6% to 38.1%. The remaining LLMs had mean scores ≥41.7% with the worst calibration observed for Meditron-7B (mean: 47.5%).

#### Molecular biology term recognition (JNLPBA)

The LLMs with the best calibration were Flan-T5-XXL (mean: 35.3%), Medicine-Llama3-8B (mean: 36.6%) and MedLLaMA-13B (mean: 39.4%). The remaining LLMs had mean scores ≥44.8%; Yi-1.5-34B-Chat had the worst calibration (mean: 57.7%).

#### Chemical recognition (BC5-chemical)

Medicine-Llama3-8B (mean: 16.5%) and Meta-Llama-3-8B-Instruct (mean: 18.8%) had the best calibration. Yi-1.5-34B, GPT-4 and GPT-3.5 had mean calibrations ranging from 20.9% to 22.9%. The remaining LLMs scored ≥40.9%.

### PICO Extraction

#### PICO extraction from abstracts (EBM-PICO)

Five LLMs had mean calibration scores ranging from 30.7% to 35.5% (ordered by increasing score: Flan-T5-XXL, Zephyr-7B-Beta, MedLLaMA-13B, Meditron-7B, Medicine-Llama3-8B) with the remaining LLMs scoring ≥49.0%.

### Relation Extraction

#### Chemical-protein relationship (ChemProt)

Flan-T5-XXL had the best calibration (mean: 32.8%), followed by Zephyr-7B-Beta (mean: 41.6%). The remaining LLMs scored ≥49.5%.

#### Drug-drug interaction (DDI)

All LLMs had a mean score ≥42.4%. The LLM with the worst calibration was Yi-1.5-34B-Chat (mean: 54.6%).

#### Gene-disease association (GAD)

Flan-T5-XXL had the best calibration (mean: 26.3%), followed by MedLLaMA-13B (mean: 31.0%) and Medicine-Llama3-8B (mean: 31.4%). The remaining LLMs all scored ≥41.7%; GPT-4 had the worst calibration (mean: 51.0%).

### Question Answering

#### Biomedical QA without context (BioASQ

Five LLMs had calibration scores between 10.3% and 14.6% (ordered by increasing score: Medicine-Llama3-8B, Meta-Llama-3-8B-Instruct, GPT-4, GPT-3.5, Yi-1.5-34B-Chat). Meditron-7B had the worst calibrtion (49.2%).

#### Biomedical QA with context (PubMedQA)

Flan-T5-XXL, Medicine-Llama3-8B, and Meta-Llama-3-8B-Instruct had mean calibration scores of 24.7%, 26.7% and 26.8% respectively. MedLLaMA-13B had a mean score of 35.7%, and the remaining LLMs had mean scores ≥41.7%.

### Sentence Similarity

#### Biomedical SS (BIOSSES)

GPT-3.5 (mean: 10.8%) and GPT-4 (mean: 13.3%) had the best calibration. Meditron-7B and Zephyr-7B-Beta had the worst calibration, scoring ≥48.6%.

### RQ1.1: Is there a relationship between calibration and family of LLMs?

The families of models (Bio-Large; Bio-Small; General-Large; General-Small) had similar mean calibrations across the tasks, ranging from 36.1% (Bio-Large) to 39.5% (General-Small) (Table 2). However, there was variation in the mean calibration scores across the families of models depending on NLP task.

**Table 2:**
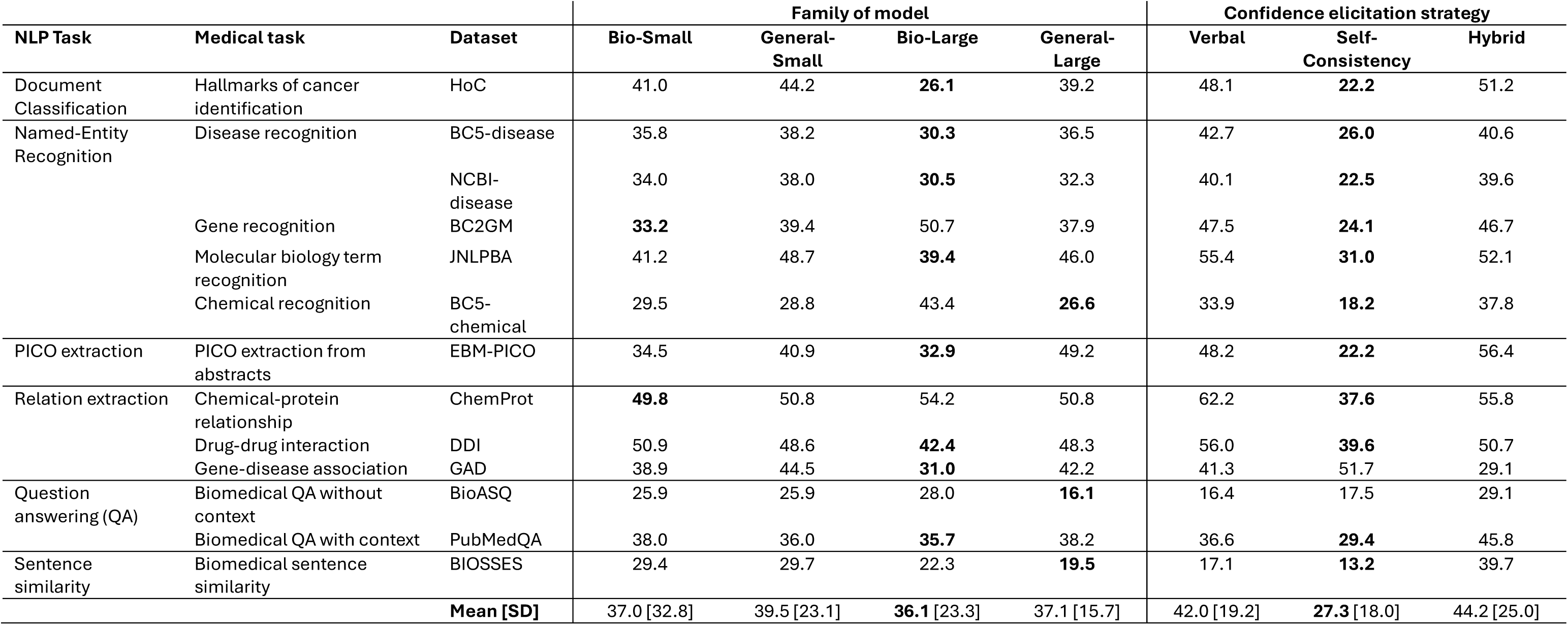
Mean Flex-ECE scores by dataset and families of models (i.e. whether LLMs are small (<10B) or large (>10B), and domain-specific (biomedical) or general domain) and confidence elicitation strategy. Only task-model-strategy combinations with a success rate ≥50% are included. Bio-small: Meditron-7B, Medicine-Llama3-8B; General-Small: Zephyr-7B-Beta, Llama-3-8B-Instruct; Bio-Large: MedLLaMa-13B; General-Large: Flan-T5-XXL, Yi-1.5-34B-Chat, GPT-3.5, GPT-4 § The calibration scores in the family of models section represent the mean of the Flex-ECE derived from each of the three confidence score elicitation strategies. PICO: Population, Intevetnion, Comparator, Outcome; QA: Question asnwering; NLP: Natural language processing; SD: Standard deviation.

The Bio-Large model (i.e. MedLLaMa-13B) had the best calibration on the Document classification task, across several of the Named-entity recognition family of tasks, and the Drug-drug interaction and Gene-disease association tasks within the Relation extraction family of tasks. However, the Bio-Large model had the worst calibration for the Chemical recognition and Gene recognition task within the Named-entity recognition family of tasks. The General-Large models had the best calibration on the Biomedical question answering task without context. The large models had better calibration than the small models on the Sentence similarity task. The families of models had relatively similar calibration across the remaining tasks.

There was also substantial variation within these families. For example, the Bio-Small family included the model with the best calibration (Medicine-Llama3-8B) and the worst calibration across the datasets (Meditron-7B).

### RQ1.2: Is there a relationship between calibration and family of BioNLP tasks?

Three families of BioNLP tasks contained more than one dataset (“Overall” rows of **Error! Reference source not found.**). Medicine-Llama3-8B had the best mean calibration scores within the Named-entity recognition family (mean: 25.7%) and the Question answering family (mean: 16.8%). Meditron-7B and Zephyr-7B-Beta had the worst calibration scores for these two BioNLP task familes; the other LLMs had broadly similar calibration scores. Regarding the Relation extraction family, the LLMs had similarly poor calibration.

### RQ1.3: Is there a relationship between calibration and confidence elicitation strategy?

Table 2Table 1 shows that the Self-consistency strategy (mean: 27.3%) was associated with the best calibration scores compared with the Verbal (mean: 42.0%) and Hybrid (mean: 44.2%) strategies across all tasks. The Self-consistency strategy had the best calibration scores for 11 of the 13 tasks, with the only exceptions being the Gene-disease association task (best calibration = hybrid strategy) and the Biomedical question answering without context (Verbal strategy had marginally better calibration).

However, this pattern was not observed for all LLMs. Four LLMs had better mean calibration scores across tasks when using the Hybrid strategy compared with the Self-consistency strategy (GPT-3.5, GPT-4, Meta-Llama-3-8B-Instruct, Yi-1.5-34B-Chat). For several models (e.g. Meditron-7B and Zephyr-7B-Beta), there was a large difference in the mean scores between the different strategies (mean score by strategy - Meditron-7B: Self-consistency = 15.6%, Hybrid = 82.3%; Zephyr-7B-Beta: Self-consistency = 24.0%, Hybrid = 66.6% [Verbal not considered for these models due to low success rate), which was less apparent for other models (e.g. Meta-Llama-3-8B-Instruct: Self-consistency = 35.7%, Hybrid = 32.4%, Verbal = 41.3%).

### RQ2: Can post-hoc calibration techniques mitigate LLM miscalibration?

When their confidences were calibrated with traditional post-hoc techniques and sufficient data, all LLMs saw substantial reduction in Flex-ECE in all tasks. Histogram binning and isotonic regression were nearly equally effective in improving calibration, reducing the average Flex-ECE across models and datasets by 23.5 percentage points (for histogram binning) and 23.6 percentage points (for isotonic regression) when using 1000 calibration examples.

As shown in Table 3, using a larger number of calibration examples consistently led to larger improvements in calibration. Though using only 10 examples for calibration decreased the mean Flex-ECE for most models, we found the pattern of improvement to be erratic, with many models seeing an increase for several datasets. We found as few as 100 examples proved sufficient to provide a large and consistent reduction in Flex-ECE, with the average Flex-ECE falling by over 75% for all models other than Medicine-Llama3-8B and MedLLaMA-13B.

**Table 3:**
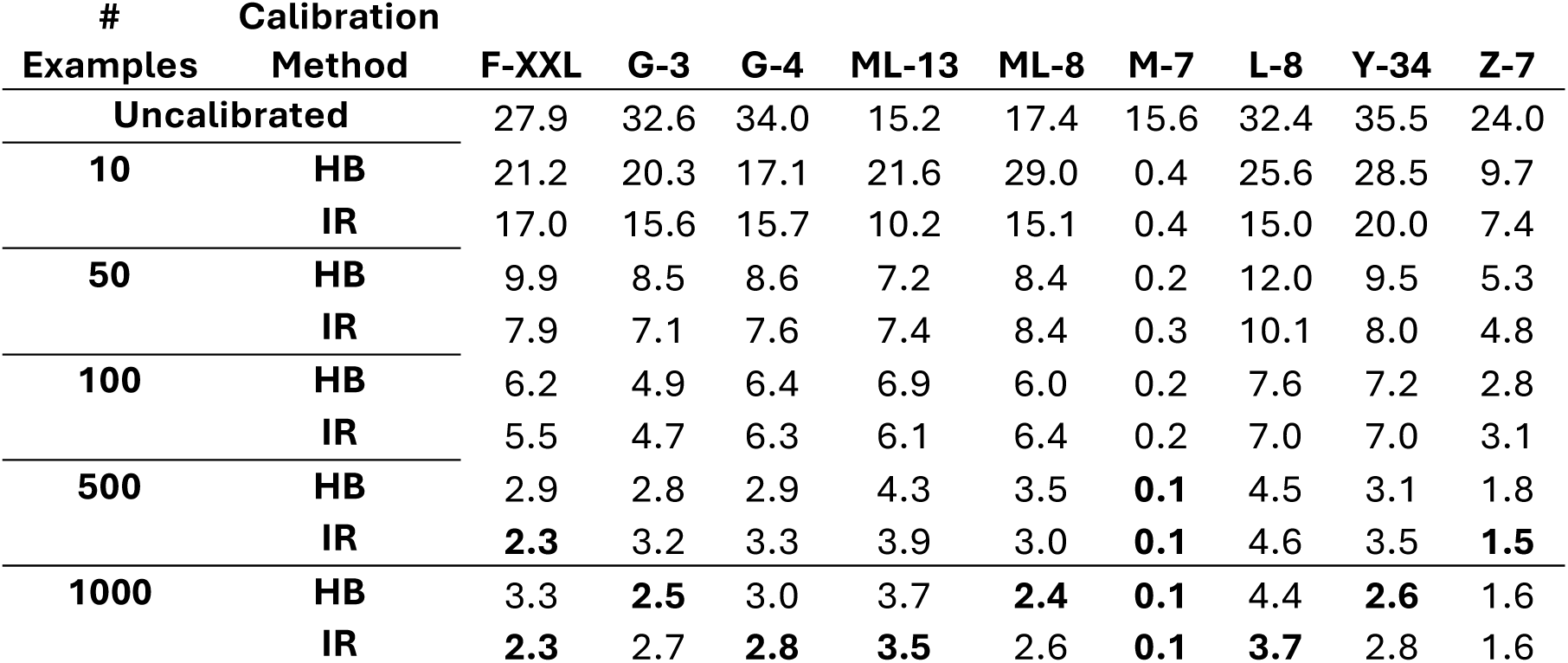
Mean calibrated Flex-ECE scores by model, maximum number of examples, and calibration method. Confidences assignment strategy used is the average best strategy for each model in RQ1. Models are abbreviated as above. Calibration methods are abbreviated: HB = Histogram binning; IR = Isotonic regression.

| # | Calibration |  |  |  |  |  |  |  |  |  |
| --- | --- | --- | --- | --- | --- | --- | --- | --- | --- | --- |
| Examples | Method | F-XXL | G-3 | G-4 | ML-13 | ML-8 | M-7 | L-8 | Y-34 | Z-7 |
| Uncalibrated |  | 27.9 | 32.6 | 34.0 | 15.2 | 17.4 | 15.6 | 32.4 | 35.5 | 24.0 |
| 10 | HB | 21.2 | 20.3 | 17.1 | 21.6 | 29.0 | 0.4 | 25.6 | 28.5 | 9.7 |
|  | IR | 17.0 | 15.6 | 15.7 | 10.2 | 15.1 | 0.4 | 15.0 | 20.0 | 7.4 |
| 50 | HB | 9.9 | 8.5 | 8.6 | 7.2 | 8.4 | 0.2 | 12.0 | 9.5 | 5.3 |
|  | IR | 7.9 | 7.1 | 7.6 | 7.4 | 8.4 | 0.3 | 10.1 | 8.0 | 4.8 |
| 100 | HB | 6.2 | 4.9 | 6.4 | 6.9 | 6.0 | 0.2 | 7.6 | 7.2 | 2.8 |
|  | IR | 5.5 | 4.7 | 6.3 | 6.1 | 6.4 | 0.2 | 7.0 | 7.0 | 3.1 |
| 500 | HB | 2.9 | 2.8 | 2.9 | 4.3 | 3.5 | 0.1 | 4.5 | 3.1 | 1.8 |
|  | IR | 2.3 | 3.2 | 3.3 | 3.9 | 3.0 | 0.1 | 4.6 | 3.5 | 1.5 |
| 1000 | HB | 3.3 | 2.5 | 3.0 | 3.7 | 2.4 | 0.1 | 4.4 | 2.6 | 1.6 |
|  | IR | 2.3 | 2.7 | 2.8 | 3.5 | 2.6 | 0.1 | 3.7 | 2.8 | 1.6 |

When a large enough number of calibration examples were used, the average Flex-ECE varied very little between tasks. For example, Flex-ECE ranged from 1.9% to 5.6% (averaged over models) for isotonic regression calibrated with a maximum of 500 examples, with QA standing out as being worse calibrated than most other tasks, having a Flex-ECE of 4.2% (though calibration for sentence similarity was technically worse, BIOSSES has far few training examples than other datasets and so cannot be easily compared).

## DISCUSSION

### Key Takeaways

#### LLM calibration is very poor in BioNLP

The main message from the results is that the current state-of-the-art methods for confidence estimation in LLMs are not trustworthy enough for real-world BioNLP solutions. The best mean Flex-ECE (Medicine-Llama3-8B, 29.8%) is not a comfortable error margin; in practice, this means that LLM confidence in BioNLP is approximately 30% off the target at best. For comparison, the worst ECE scores in [1] also for NLP tasks, albeit non-BioNLP, was approximately 8%, far below the best mean Flex-ECE we found in this BioNLP-specific study.

#### LLMs tend to be over-confident

The models exhibited general overconfidence (see reliability diagrams in Supplement File 3) in their answers when questioned. This inability to exhibit uncertainty is a major blocker to the trustworthiness and therefore usability of these models, regardless of accuracy.

#### No ‘one size fits all’ approach

We could not establish many reliable patterns between calibration and LLM families or confidence assignment strategies, which suggests that a tailored approach to assigning confidence scores to LLM responses is required in real-world, biomedical applications. The only clear pattern we observed is that self-consistency was consistently more reliable than other strategies (lowest mean Flex-ECE in 11/13 tasks).

#### Post-hoc calibration helps

As a result, users are warned to place the necessary *guardrails* around LLMs when building LLM-backed, BioNLP solutions that rely on confidence estimation. Traditional post-hoc calibration techniques proved promising: despite being low-cost, they drastically reduced calibration errors, thereby increasing the degree of reliability of the confidence estimations we investigated in this study.

### Known Limitations

#### Verbal confidence elicitation

Because LLMs are highly susceptible to the wordings of prompts [22–24] it may be that the prompts to elicit verbal confidence estimations can be improved. Techniques such as *Chain-of-Thought* [25] and *p-tuning* [26] have proven useful to increase both LLM performance and reliability.

#### Verbal confidence parsing

Because LLMs can output any text, including unintelligible *gibberish*, verbal confidence estimation is often not feasible in an automated framework for confidence estimation, evaluation and mitigation such as ours.

Techniques such as *declarative prompting* [27,28] that constraint LLM output may help avoid such scenarios.

#### Calibration is not utility

ECE (flex or not) can be a misleading quantification of reliability, because models can be very well calibrated, yet have poor performance, i.e. low utility. This is particularly true when confidence scores cluster around a certain range (i.e. a *bucket*), a phenomenon dubbed *bucket collapse* [29]. We observed this mostly with the calibrated scores, i.e. after-post-hoc calibration, when we ended up with most confidence scores *clumped* around the same low range of the LLM’s performance, so that calibration could be improved. We urge users to couple (Flex-)ECE with the usual performance metrics (e.g. F1 or accuracy) when judging the reliability of LLMs (and confidence estimation strategies).

#### Black-box confidence

All calibration and confidence estimation methods used in this study are designed to work with *black-box* access to an LLM, i.e. users need not have access to the inner workings of LLMs to quantify and/or mitigate miscalibration. This was a deliberate decision in this study to allow any user to evaluate any LLM and/or apply this framework in their own work. Advanced users may wish to explore white-box confidence estimation techniques, such as *aggregating token probabilities* [30], as they have shown some degree of success. Note that black-box use of closed-source LLMs such as GPT is the only way to use such LLMs by any user, advanced or not.

## CONCLUSION

This study assesses the calibration of LLMs in BioNLP. Results corroborate previous studies that neural networks are poorly calibrated but highlights an even more alarming situation when networks are LLMs and the tasks are biomedically related: at best, LLMs are approximately 30% off the target in the scenarios we investigated.

In addition to the above findings, we also introduced Flex-ECE, a calibration metric that more realistically quantifies calibration of NLP models, and indeed any model that can be partially right or wrong. Flex-ECE is a key contribution of this study to the scientific community, as it opens path for future work to explore calibration methods that move away from binary correctness and towards a more continuous notion of correctness.

## Footnotes

1 https://microsoft.github.io/BLURB/tasks.html#overview

2 Prompt templates for all datasets in Supplemental File 1.

3 Modern LLMs operate with “tokens” rather than “words”, where a “token” can be a whole word or just part of a word. For instance, it could be that an LLM models the word “apple” as 1 token but “misplace” as 2 tokens: “mis” + “place”. Tokens can also be non-words such as “new line” or “end of generation”.

4 Confidence assignment strategies formally defined in Supplemental File 2.

5 To guarantee that the LLM *can* vary responses, we prompt the LLM with an increasing degree of freedom when sampling, by increasing the parameter *temperature* before each call.

6 Flex-ECE formally defined in Supplemental File 2.

7 Similarity functions formally defined in Supplemental File 2.

## REFERENCES

1 Guo C, Pleiss G, Sun Y, et al. On Calibration of Modern Neural Networks. In: Precup D, Teh YW, eds. Proceedings of the 34th International Conference on Machine Learning. PMLR 2017:1321–30.

2 Xiong M, Hu Z, Lu X, et al. Can LLMs Express Their Uncertainty? An Empirical Evaluation of Confidence Elicitation in LLMs. The Twelfth International Conference on Learning Representations. 2024.

3 Liu Y, He H, Han T, et al. Understanding LLMs: A Comprehensive Overview from Training to Inference. 2024.

4 Huang Y, Sun L, Wang H, et al. TrustLLM: Trustworthiness in Large Language Models. 2024.

5 Geng J, Cai F, Wang Y, et al. A Survey of Confidence Estimation and Calibration in Large Language Models. In: Duh K, Gomez H, Bethard S, eds. Proceedings of the 2024 Conference of the North American Chapter of the Association for Computational Linguistics: Human Language Technologies (Volume 1: Long Papers). Mexico City, Mexico: Association for Computational Linguistics 2024:6577–95.

6 Feng H, Ronzano F, LaFleur J, et al. Evaluation of large language model performance on the Biomedical Language Understanding and Reasoning Benchmark. 2024.

7 Tunstall L, Beeching E, Lambert N, et al. Zephyr: Direct Distillation of LM Alignment. 2023.

8 AI@Meta. Llama 3 Model Card. Published Online First: 2024.

9 Chung HW, Hou L, Longpre S, et al. Scaling Instruction-Finetuned Language Models. Journal of Machine Learning Research. 2024;25:1–53.

10 AI 01, Young A, Chen B, et al. Yi: Open Foundation Models by 01.AI. 2024.

11 Azure OpenAI Service Models. https://learn.microsoft.com/en-us/azure/ai-services/openai/concepts/models (accessed 17 December 2024)

12 Chen Z, Hernández-Cano A, Romanou A, et al. MEDITRON-70B: Scaling Medical Pretraining for Large Language Models. 2023.

13 Cheng D, Gu Y, Huang S, et al. Instruction Pre-Training: Language Models are Supervised Multitask Learners. arXiv preprint arXiv:240614491. 2024.

14 Cheng D, Huang S, Wei F. Adapting Large Language Models via Reading Comprehension. The Twelfth International Conference on Learning Representations. 2024.

15 Wu C, Lin W, Zhang X, et al. PMC-LLaMA: toward building open-source language models for medicine. Journal of the American Medical Informatics Association. 2024;31:1833–43. doi: 10.1093/jamia/ocae045

16 Tian K, Mitchell E, Zhou A, et al. Just Ask for Calibration: Strategies for Eliciting Calibrated Confidence Scores from Language Models Fine-Tuned with Human Feedback. The 2023 Conference on Empirical Methods in Natural Language Processing. 2023.

17 Huang Y, Liu Y, Thirukovalluru R, et al. Calibrating Long-form Generations from Large Language Models. 2024.

18 Xia Y, Kim J, Chen Y, et al. Understanding the Performance and Estimating the Cost of LLM Fine-Tuning. 2024 IEEE International Symposium on Workload Characterization (IISWC). 2024:210–23.

19 Zadrozny B, Elkan C. Transforming classifier scores into accurate multiclass probability estimates. Proceedings of the eighth ACM SIGKDD international conference on Knowledge discovery and data mining. 2002:694–9.

20 Zadrozny B, Elkan C. Obtaining calibrated probability estimates from decision trees and naive bayesian classifiers. Icml. 2001:609–16.

21 Niculescu-Mizil A, Caruana R. Predicting good probabilities with supervised learning. Proceedings of the 22nd international conference on Machine learning - ICML’05. Bonn, Germany: ACM Press 2005:625–32.

22 Sclar M, Choi Y, Tsvetkov Y, et al. Quantifying Language Models’ Sensitivity to Spurious Features in Prompt Design or: How I learned to start worrying about prompt formatting. The Twelfth International Conference on Learning Representations. 2024.

23 Zhuo J, Zhang S, Fang X, et al. ProSA: Assessing and Understanding the Prompt Sensitivity of LLMs. In: Al-Onaizan Y, Bansal M, Chen Y-N, eds. Findings of the Association for Computational Linguistics: EMNLP 2024. Miami, Florida, USA: Association for Computational Linguistics 2024:1950–76.

24 Anagnostidis S, Bulian J. How Susceptible are LLMs to Influence in Prompts? First Conference on Language Modeling. 2024.

25 Wei J, Wang X, Schuurmans D, et al. Chain-of-Thought Prompting Elicits Reasoning in Large Language Models. In: Koyejo S, Mohamed S, Agarwal A, et al., eds. Advances in Neural Information Processing Systems. Curran Associates, Inc. 2022:24824–37.

26 Lester B, Al-Rfou R, Constant N. The Power of Scale for Parameter-Efficient Prompt Tuning. In: Moens M-F, Huang X, Specia L, et al., eds. Proceedings of the 2021 Conference on Empirical Methods in Natural Language Processing. Online and Punta Cana, Dominican Republic: Association for Computational Linguistics 2021:3045–59.

27 Willard BT, Louf R. Efficient Guided Generation for Large Language Models. 2023.

28 Khattab O, Singhvi A, Maheshwari P, et al. DSPy: Compiling Declarative Language Model Calls into Self-Improving Pipelines. The Twelfth International Conference on Learning Representations. 2024.

29 Spiess C, Gros D, Pai KS, et al. Calibration and Correctness of Language Models for Code. 2024.

30 Kumar A, Morabito R, Umbet S, et al. Confidence Under the Hood: An Investigation into the Confidence-Probability Alignment in Large Language Models. 2024.

