## Supplemental File 1 for "A Study of Calibration as a Measurement of Trustworthiness of Large Language Models in Biomedical Research"

### Supplementary File 1 – Prompt Templates

This is a supplementary file to a full manuscript *A Study of Calibration as a Measurement of Trustworthiness of Large Language Models in Biomedical Research*.

#### Prompt Templates

##### Structure & Verbosity

Although the content of each prompt template was the same for each dataset, the *style* of each template differed in terms of *verbosity* and *structure*. The verbosity—short vs long—was the best-performing template verbosity as in Feng et al, 2024, while the structure is determined by the type of large language model (LLM).

The below table informs which prompt *structure* different LLMs require:

| Basic | Chat |
|---|---|
| • Flan-T5-XXL<br>• Medicine-Llama3-8B<br>• Meditron-7B<br>• MedLLaMA-13B | • GPT-3.5-Turbo<br>• GPT-4<br>• Llama-3-8B-Instruct<br>• Yi-1.5-34B-Chat<br>• Zephyr-7B-Beta |

The below table shows the selected *verbosity* for each model-dataset pair:

| LLM | Short | Long |
|---|---|---|
| **Flan-T5-XXL** | BC2GM, BC5-chemical, BC5-disease, JNLPBA, NCBI-disease, EBM-PICO, GAD, BIOSSES, HoC | DDI, ChemProt, BioASQ, PubMedQA |
| **GPT-3.5** | BC2GM, BC5-disease, JNLPBA, EBM-PICO, GAD, HoC, BioASQ, PubMedQA | BC5-chemical, NCBI-disease, DDI, ChemProt, BIOSSES |
| **GPT-4** | BC2GM, BC5-disease, JNLPBA, NCBI-disease, EBM-PICO, DDI, BIOSSES, HoC | BC5-chemical, ChemProt, GAD, BioASQ, PubMedQA |
| **Meta-Llama-3-8B-Instruct** | EBM-PICO, BIOSSES, HoC, BioASQ | BC2GM, BC5-chemical, BC5-disease, JNLPBA, NCBI-disease, DDI, ChemProt, GAD, PubMedQA |
| **Medicine-Llama3-8B** | BIOSSES | BC2GM, BC5-chemical, BC5-disease, JNLPBA, NCBI-disease, EBM-PICO, DDI, ChemProt, GAD, HoC, BioASQ, PubMedQA |

| | | |
|---|---|---|
| **Meditron-7B** | BC2GM, JNLPBA, BIOSSES, PubMedQA | BC5-chemical, BC5-disease, NCBI-disease, EBM-PICO, DDI, ChemProt, GAD, HoC, BioASQ |
| **MedLLaMA-13B** | JNLPBA, EBM-PICO, PubMedQA | BC2GM, BC5-chemical, BC5-disease, NCBI-disease, DDI, ChemProt, GAD, BIOSSES, HoC, BioASQ |
| **Yi-1.5-34B-Chat** | BC2GM, BC5-chemical, DDI, GAD, HoC | BC5-disease, JNLPBA, NCBI-disease, EBM-PICO, ChemProt, BIOSSES, BioASQ, PubMedQA |
| **Zephyr-7B-Beta** | BC2GM, EBM-PICO, GAD, HoC, BioASQ | BC5-chemical, BC5-disease, JNLPBA, NCBI-disease, DDI, ChemProt, BIOSSES, PubMedQA |

#### Templates

The below tables show the prompt template set for each dataset. At runtime, the final prompt to the fired to the LLM is composed by replacing placeholders *{text}* (or *{sent-1}* and *{sent-2}* in the case of HoC) with the actual example in the dataset.

##### DC – HoC

| Style | Template |
|---|---|
| **Short, basic** | ***INSTRUCTIONS***<br>You are an expert in the biomedical domain. You are a smart document classification assistant, specialized in biomedical document classification. Classify the provided abstract. If the abstract does not belong to any of the hallmarks of cancer, answer with EMPTY_LIST. Otherwise, answer with a comma-separated list consisting of one or multiple of the following hallmarks of cancer:<br>activating invasion and metastasis<br>sustaining proliferative signaling<br>resisting cell death<br>cellular energetics<br>genomic instability and mutation<br>evading growth suppressors<br>inducing angiogenesis<br>enabling replicative immortality<br>avoiding immune destruction<br>tumor promoting inflammation<br><br>***YOUR TURN***<br>INPUT: sentence1: {sent-1} sentence2: {sent-2}<br>OUTPUT: |
| **Long, basic** | ***INSTRUCTIONS***<br>You are an expert in the biomedical domain. You are a smart |

| | |
|---|---|
| | document classification assistant, specialized in biomedical document classification. I will provide you the output format, the definition of class categories, and the abstract to classify.<br>Answer with a comma-separated list consisting of one or multiple hallmarks of cancer.<br>Include the class in the list if the abstract is related to that class.<br>Please note one abstract can be related to multiple classes.<br>If the abstract does not belong to any of the hallmarks of cancer, answer with EMPTY_LIST<br>Output Format:<br>Category Definition:<br>There are 10 cancer hallmarks you will need to decide whether the article is related to, including:<br>activating invasion and metastasis<br>sustaining proliferative signaling<br>resisting cell death<br>cellular energetics<br>genomic instability and mutation<br>evading growth suppressors<br>inducing angiogenesis<br>enabling replicative immortality<br>avoiding immune destruction<br>tumor promoting inflammation<br><br>***YOUR TURN***<br>INPUT: sentence1: {sent-1} sentence2: {sent-2}<br>OUTPUT: |
| **Short, chat** | SYSTEM: You are an expert in the biomedical domain. You are a smart document classification assistant, specialized in biomedical document classification. Classify the provided abstract. If the abstract does not belong to any of the hallmarks of cancer, answer with EMPTY_LIST. Otherwise, answer with a comma-separated list consisting of one or multiple of the following hallmarks of cancer:<br>activating invasion and metastasis<br>sustaining proliferative signaling<br>resisting cell death<br>cellular energetics<br>genomic instability and mutation<br>evading growth suppressors<br>inducing angiogenesis<br>enabling replicative immortality<br>avoiding immune destruction<br>tumor promoting inflammation<br><br>USER: sentence1: {sent-1} sentence2: {sent-2} |
| **Long, chat** | SYSTEM: You are an expert in the biomedical domain. You are a smart document classification assistant, specialized in biomedical |

| | |
|---|---|
| | document classification. I will provide you the output format, the definition of class categories, and the abstract to classify.<br>Answer with a comma-separated list consisting of one or multiple hallmarks of cancer.<br>Include the class in the list if the abstract is related to that class.<br>Please note one abstract can be related to multiple classes.<br>If the abstract does not belong to any of the hallmarks of cancer, answer with EMPTY_LIST<br>Output Format:<br>Category Definition:<br>There are 10 cancer hallmarks you will need to decide whether the article is related to, including:<br>activating invasion and metastasis<br>sustaining proliferative signaling<br>resisting cell death<br>cellular energetics<br>genomic instability and mutation<br>evading growth suppressors<br>inducing angiogenesis<br>enabling replicative immortality<br>avoiding immune destruction<br>tumor promoting inflammation<br><br>USER: sentence1: {sent-1} sentence2: {sent-2} |

#### NER – BC2GM

| Style | Template |
|---|---|
| **Short, basic** | ***INSTRUCTIONS***<br>You are an expert in the biomedical domain. You are a smart named entity recognition assistant, specialized in extracting mentions of gene entities from text. Based on the provided sentence, answer with NO_ENTITIES if you find no mentions of gene entities, otherwise, answer with a comma-delimited list of the mentions of gene entities you find. Do not provide any explanation or other characters.<br><br>***YOUR TURN***<br>INPUT: {text}<br>OUTPUT: |
| **Long, basic** | ***INSTRUCTIONS***<br>You are an expert in the biomedical domain. You are a smart named entity recognition assistant, specialized in extracting mentions of gene entities from text. I will provide you the output format, the definition of the entities you need to extract, and the sentence from which to extract the entities.<br>Answer with a comma-separated list of gene entities. If no gene entities are present in the sentence, answer with NO_ENTITIES. Do not |

| | |
|---|---|
| | provide any explanation or other characters.<br>Output Format:<br>gene: a basic unit of heredity and a sequence of nucleotides in DNA.<br><br>***YOUR TURN***<br>INPUT: {text}<br>OUTPUT: |
| **Short, chat** | SYSTEM: You are an expert in the biomedical domain. You are a smart named entity recognition assistant, specialized in extracting mentions of gene entities from text. Based on the provided sentence, answer with NO_ENTITIES if you find no mentions of gene entities, otherwise, answer with a comma-delimited list of the mentions of gene entities you find. Do not provide any explanation or other characters.<br><br>USER: {text} |
| **Long, chat** | SYSTEM: You are an expert in the biomedical domain. You are a smart named entity recognition assistant, specialized in extracting mentions of gene entities from text. I will provide you the output format, the definition of the entities you need to extract, and the sentence from which to extract the entities.<br>Answer with a comma-separated list of gene entities. If no gene entities are present in the sentence, answer with NO_ENTITIES. Do not provide any explanation or other characters.<br>Output Format:<br>gene: a basic unit of heredity and a sequence of nucleotides in DNA.<br><br>USER: {text} |

#### NER – BC5-Chemical

| Style | Template |
|---|---|
| **Short, basic** | ***INSTRUCTIONS***<br>You are an expert in the biomedical domain. You are a smart named entity recognition assistant, specialized in extracting mentions of chemical entities from text. Based on the provided sentence, answer with NO_ENTITIES if you find no mentions of chemical entities, otherwise, answer with a comma-delimited list of the mentions of chemical entities you find. Do not provide any explanation or other characters.<br><br>***YOUR TURN***<br>INPUT: {text}<br>OUTPUT: |
| **Long, basic** | ***INSTRUCTIONS***<br>You are an expert in the biomedical domain. You are a smart named entity recognition assistant, specialized in extracting mentions of chemical entities from text. I will provide you the output format, the definition of the entities you need to extract, and the sentence from |

| | |
|---|---|
| | which to extract the entities.<br>Answer with a comma-separated list of chemical entities. If no chemical entities are present in the sentence, answer with NO_ENTITIES. Do not provide any explanation or other characters.<br>Output Format:<br>chemical: a compound or substance that has been purified or prepared.<br><br>***YOUR TURN***<br>INPUT: {text}<br>OUTPUT: |
| **Short, chat** | SYSTEM: You are an expert in the biomedical domain. You are a smart named entity recognition assistant, specialized in extracting mentions of chemical entities from text. Based on the provided sentence, answer with NO_ENTITIES if you find no mentions of chemical entities, otherwise, answer with a comma-delimited list of the mentions of chemical entities you find. Do not provide any explanation or other characters.<br><br>USER: {text} |
| **Long, chat** | SYSTEM: You are an expert in the biomedical domain. You are a smart named entity recognition assistant, specialized in extracting mentions of chemical entities from text. I will provide you the output format, the definition of the entities you need to extract, and the sentence from which to extract the entities.<br>Answer with a comma-separated list of chemical entities. If no chemical entities are present in the sentence, answer with NO_ENTITIES. Do not provide any explanation or other characters.<br>Output Format:<br>chemical: a compound or substance that has been purified or prepared.<br><br>USER: {text} |

#### NER – BC5-Disease

| **Style** | **Template** |
|---|---|
| **Short, basic** | ***INSTRUCTIONS***<br>You are an expert in the biomedical domain. You are a smart named entity recognition assistant, specialized in extracting mentions of disease entities from text. Based on the provided sentence, answer with NO_ENTITIES if you find no mentions of disease entities, otherwise, answer with a comma-delimited list of the mentions of disease entities you find. Do not provide any explanation or other characters.<br><br>***YOUR TURN*** |

| | |
|---|---|
| | INPUT: {text}<br>OUTPUT: |
| **Long, basic** | ***INSTRUCTIONS***<br>You are an expert in the biomedical domain. You are a smart named entity recognition assistant, specialized in extracting mentions of disease entities from text. I will provide you the output format, the definition of the entities you need to extract, and the sentence from which to extract the entities.<br>Answer with a comma-separated list of disease entities. If no disease entities are present in the sentence, answer with NO_ENTITIES. Do not provide any explanation or other characters.<br>Output Format:<br>disease: a disorder of structure or function in a human, animal, or plant, especially one that has a known cause and a distinctive group of symptoms, signs, or anatomical changes.<br><br>***YOUR TURN***<br>INPUT: {text}<br>OUTPUT: |
| **Short, chat** | SYSTEM: You are an expert in the biomedical domain. You are a smart named entity recognition assistant, specialized in extracting mentions of disease entities from text. Based on the provided sentence, answer with NO_ENTITIES if you find no mentions of disease entities, otherwise, answer with a comma-delimited list of the mentions of disease entities you find. Do not provide any explanation or other characters.<br><br>USER: {text} |
| **Long, chat** | SYSTEM: You are an expert in the biomedical domain. You are a smart named entity recognition assistant, specialized in extracting mentions of disease entities from text. I will provide you the output format, the definition of the entities you need to extract, and the sentence from which to extract the entities.<br>Answer with a comma-separated list of disease entities. If no disease entities are present in the sentence, answer with NO_ENTITIES. Do not provide any explanation or other characters.<br>Output Format:<br>disease: a disorder of structure or function in a human, animal, or plant, especially one that has a known cause and a distinctive group of symptoms, signs, or anatomical changes.<br><br>USER: {text} |

#### NER – JNLPBA

| Style | Template |
|---|---|

| | |
|---|---|
| **Short, basic** | ***INSTRUCTIONS***<br>You are an expert in the biomedical domain. You are a smart named entity recognition assistant, specialized in extracting mentions of protein, gene, and cell entities from text. Based on the provided sentence, answer with NO_ENTITIES if you find no mentions of protein, gene, or cell entities, otherwise, answer with a comma-delimited list of the mentions of protein, gene, or cell entities you find. Do not provide any explanation or other characters.<br><br>***YOUR TURN***<br>INPUT: {text}<br>OUTPUT: |
| **Long, basic** | ***INSTRUCTIONS***<br>You are an expert in the biomedical domain. You are a smart named entity recognition assistant, specialized in extracting mentions of protein, gene, and cell entities from text. I will provide you the output format, the definition of the entities you need to extract, and the sentence from which to extract the entities.<br>Answer with a comma-separated list of protein, gene, and cell entities. If no protein, gene, or cell entities are present in the sentence, answer with NO_ENTITIES. Do not provide any explanation or other characters.<br>Output Format:<br>cell: includes cell line (a cell culture developed from a single cell and therefore consisting of cells with a uniform genetic makeup) and cell type (a classification used to identify cells that share morphological or phenotypical features)<br>gene: a basic unit of heredity and a sequence of nucleotides in DNA.<br>protein: a large, complex molecule that play many critical roles in a living organism.<br><br>***YOUR TURN***<br>INPUT: {text}<br>OUTPUT: |
| **Short, chat** | SYSTEM: You are an expert in the biomedical domain. You are a smart named entity recognition assistant, specialized in extracting mentions of protein, gene, and cell entities from text. Based on the provided sentence, answer with NO_ENTITIES if you find no mentions of protein, gene, or cell entities, otherwise, answer with a comma-delimited list of the mentions of protein, gene, or cell entities you find. Do not provide any explanation or other characters.<br><br>USER: {text} |
| **Long, chat** | SYSTEM: You are an expert in the biomedical domain. You are a smart named entity recognition assistant, specialized in extracting mentions of protein, gene, and cell entities from text. I will provide you the output format, the definition of the entities you need to extract, and the sentence from which to extract the entities. |

| | |
|---|---|
| | Answer with a comma-separated list of protein, gene, and cell entities. If no protein, gene, or cell entities are present in the sentence, answer with NO_ENTITIES. Do not provide any explanation or other characters.<br>Output Format:<br>cell: includes cell line (a cell culture developed from a single cell and therefore consisting of cells with a uniform genetic makeup) and cell type (a classification used to identify cells that share morphological or phenotypical features)<br>gene: a basic unit of heredity and a sequence of nucleotides in DNA.<br>protein: a large, complex molecule that play many critical roles in a living organism.<br><br>USER: {text} |

##### NER – NCBI-Disease

| Style | Template |
|---|---|
| **Short, basic** | ***INSTRUCTIONS***<br>You are an expert in the biomedical domain. You are a smart named entity recognition assistant, specialized in extracting mentions of disease entities from text. Based on the provided sentence, answer with NO_ENTITIES if you find no mentions of disease entities, otherwise, answer with a comma-delimited list of the mentions of disease entities you find. Do not provide any explanation or other characters.<br><br>***YOUR TURN***<br>INPUT: {text}<br>OUTPUT: |
| **Long, basic** | ***INSTRUCTIONS***<br>You are an expert in the biomedical domain. You are a smart named entity recognition assistant, specialized in extracting mentions of disease entities from text. I will provide you the output format, the definition of the entities you need to extract, and the sentence from which to extract the entities.<br>Answer with a comma-separated list of disease entities. If no disease entities are present in the sentence, answer with NO_ENTITIES. Do not provide any explanation or other characters.<br>Output Format:<br>disease: a disorder of structure or function in a human, animal, or plant, especially one that has a known cause and a distinctive group of symptoms, signs, or anatomical changes.<br><br>***YOUR TURN***<br>INPUT: {text}<br>OUTPUT: |
| **Short, chat** | SYSTEM: You are an expert in the biomedical domain. You are a smart named entity recognition assistant, specialized in extracting mentions |

| | |
|---|---|
| | of disease entities from text. Based on the provided sentence, answer with NO_ENTITIES if you find no mentions of disease entities, otherwise, answer with a comma-delimited list of the mentions of disease entities you find. Do not provide any explanation or other characters.<br><br>USER: {text} |
| **Long, chat** | SYSTEM: You are an expert in the biomedical domain. You are a smart named entity recognition assistant, specialized in extracting mentions of disease entities from text. I will provide you the output format, the definition of the entities you need to extract, and the sentence from which to extract the entities.<br>Answer with a comma-separated list of disease entities. If no disease entities are present in the sentence, answer with NO_ENTITIES. Do not provide any explanation or other characters.<br>Output Format:<br>disease: a disorder of structure or function in a human, animal, or plant, especially one that has a known cause and a distinctive group of symptoms, signs, or anatomical changes.<br><br>USER: {text} |

#### PICO – EBM-PICO – Intervention

| Style | Template |
|---|---|
| **Short, basic** | ***INSTRUCTIONS***<br>You are an expert in the biomedical domain. You are a smart named entity recognition assistant, specialized in extracting mentions of interventions of clinical trials from text. Based on the provided clinical trial report, answer with NO_ENTITIES if you find no mentions of clinical trial interventions, otherwise, answer with a comma-delimited list of the mentions of clinical trial interventions you find. Do not provide any explanation or other characters.<br><br>***YOUR TURN***<br>INPUT: {text}<br>OUTPUT: |
| **Long, basic** | ***INSTRUCTIONS***<br>You are an expert in the biomedical domain. You are a smart named entity recognition assistant, specialized in extracting mentions of interventions of clinical trials from text. I will provide you the output format, the definition of the entities you need to extract, and the sentence from which to extract the entities.<br>Answer with a comma-separated list of clinical trial interventions. If no interventions are present in the sentence, answer with NO_ENTITIES. Do not provide any explanation or other characters.<br>Output Format: |

| | |
|---|---|
| | clinical trial intervention: a potential drug, medical device, activity, or procedure.<br><br>***YOUR TURN***<br>INPUT: {text}<br>OUTPUT: |
| **Short, chat** | SYSTEM: You are an expert in the biomedical domain. You are a smart named entity recognition assistant, specialized in extracting mentions of interventions of clinical trials from text. Based on the provided clinical trial report, answer with NO_ENTITIES if you find no mentions of clinical trial interventions, otherwise, answer with a comma-delimited list of the mentions of clinical trial interventions you find. Do not provide any explanation or other characters.<br><br>USER: {text} |
| **Long, chat** | SYSTEM: You are an expert in the biomedical domain. You are a smart named entity recognition assistant, specialized in extracting mentions of interventions of clinical trials from text. I will provide you the output format, the definition of the entities you need to extract, and the sentence from which to extract the entities.<br>Answer with a comma-separated list of clinical trial interventions. If no interventions are present in the sentence, answer with NO_ENTITIES. Do not provide any explanation or other characters.<br>Output Format:<br>clinical trial intervention: a potential drug, medical device, activity, or procedure.<br><br>USER: {text} |

#### PICO – EBM-PICO – Outcome

| Style | Template |
|---|---|
| **Short, basic** | ***INSTRUCTIONS***<br>You are an expert in the biomedical domain. You are a smart named entity recognition assistant, specialized in extracting mentions of outcomes of clinical trials from text. Based on the provided clinical trial report, answer with NO_ENTITIES if you find no mentions of clinical trial outcomes, otherwise, answer with a comma-delimited list of the mentions of clinical trial outcomes you find. Do not provide any explanation or other characters.<br><br>***YOUR TURN***<br>INPUT: {text}<br>OUTPUT: |
| **Long, basic** | ***INSTRUCTIONS***<br>You are an expert in the biomedical domain. You are a smart named entity recognition assistant, specialized in extracting mentions of |

| | |
|---|---|
| | outcomes of clinical trials from text. I will provide you the output format, the definition of the entities you need to extract, and the sentence from which to extract the entities.<br>Answer with a comma-separated list of clinical trial outcomes. If no outcomes are present in the sentence, answer with NO_ENTITIES. Do not provide any explanation or other characters.<br>Output Format:<br>clinical trial outcome: the impact that a given intervention or exposure has on the health of clinical trial participants.<br><br>***YOUR TURN***<br>INPUT: {text}<br>OUTPUT: |
| **Short, chat** | SYSTEM: You are an expert in the biomedical domain. You are a smart named entity recognition assistant, specialized in extracting mentions of outcomes of clinical trials from text. Based on the provided clinical trial report, answer with NO_ENTITIES if you find no mentions of clinical trial outcomes, otherwise, answer with a comma-delimited list of the mentions of clinical trial outcomes you find. Do not provide any explanation or other characters.<br><br>USER: {text} |
| **Long, chat** | SYSTEM: You are an expert in the biomedical domain. You are a smart named entity recognition assistant, specialized in extracting mentions of outcomes of clinical trials from text. I will provide you the output format, the definition of the entities you need to extract, and the sentence from which to extract the entities.<br>Answer with a comma-separated list of clinical trial outcomes. If no outcomes are present in the sentence, answer with NO_ENTITIES. Do not provide any explanation or other characters.<br>Output Format:<br>clinical trial outcome: the impact that a given intervention or exposure has on the health of clinical trial participants.<br><br>USER: {text} |

#### PICO – EBM-PICO – Participant

| Style | Template |
|---|---|
| **Short, basic** | ***INSTRUCTIONS***<br>You are an expert in the biomedical domain. You are a smart named entity recognition assistant, specialized in extracting mentions of participants of clinical trials from text. Based on the provided clinical trial report, answer with NO_ENTITIES if you find no mentions of clinical trial participants, otherwise, answer with a comma-delimited list of the mentions of clinical trial participants you find. Do not provide any explanation or other characters. |

| | |
|---|---|
| | ***YOUR TURN***<br>INPUT: {text}<br>OUTPUT: |
| **Long, basic** | ***INSTRUCTIONS***<br>You are an expert in the biomedical domain. You are a smart named entity recognition assistant, specialized in extracting mentions of participants of clinical trials from text. I will provide you the output format, the definition of the entities you need to extract, and the sentence from which to extract the entities.<br>Answer with a comma-separated list of clinical trial participants. If no participants are present in the sentence, answer with NO_ENTITIES. Do not provide any explanation or other characters.<br>Output Format:<br>clinical trial participant: a person who takes part in a clinical trial.<br><br>***YOUR TURN***<br>INPUT: {text}<br>OUTPUT: |
| **Short, chat** | SYSTEM: You are an expert in the biomedical domain. You are a smart named entity recognition assistant, specialized in extracting mentions of participants of clinical trials from text. Based on the provided clinical trial report, answer with NO_ENTITIES if you find no mentions of clinical trial participants, otherwise, answer with a comma-delimited list of the mentions of clinical trial participants you find. Do not provide any explanation or other characters.<br><br>USER: {text} |
| **Long, chat** | SYSTEM: You are an expert in the biomedical domain. You are a smart named entity recognition assistant, specialized in extracting mentions of participants of clinical trials from text. I will provide you the output format, the definition of the entities you need to extract, and the sentence from which to extract the entities.<br>Answer with a comma-separated list of clinical trial participants. If no participants are present in the sentence, answer with NO_ENTITIES. Do not provide any explanation or other characters.<br>Output Format:<br>clinical trial participant: a person who takes part in a clinical trial.<br><br>USER: {text} |

#### QA – BioASQ

| Style | Template |
|---|---|
| **Short, basic** | ***INSTRUCTIONS***<br>You are an expert in the biomedical domain. You are a smart question answering assistant, specialized in answering biomedical questions. Only answer with yes, or no. Do not provide any explanation or other |

| | |
|---|---|
| | characters.<br><br>***YOUR TURN***<br>INPUT: {text}<br>OUTPUT: |
| **Long, basic** | ***INSTRUCTIONS***<br>You are an expert in the biomedical domain. You are a smart question answering assistant, specialized in answering biomedical questions. I will provide you the output format, and the question to answer.<br>Your answer must be one word: yes, or no. Do not provide any explanation or other characters.<br><br>***YOUR TURN***<br>INPUT: {text}<br>OUTPUT: |
| **Short, chat** | SYSTEM: You are an expert in the biomedical domain. You are a smart question answering assistant, specialized in answering biomedical questions. Only answer with yes, or no. Do not provide any explanation or other characters.<br><br>USER: {text} |
| **Long, chat** | SYSTEM: You are an expert in the biomedical domain. You are a smart question answering assistant, specialized in answering biomedical questions. I will provide you the output format, and the question to answer.<br>Your answer must be one word: yes, or no. Do not provide any explanation or other characters.<br><br>USER: {text} |

#### QA – PubMedQA

| Style | Template |
|---|---|
| **Short, basic** | ***INSTRUCTIONS***<br>You are an expert in the biomedical domain. You are a smart question answering assistant, specialized in answering biomedical questions. Answer based on the provided abstract. Only answer with yes, no, or maybe. Do not provide any explanation or other characters.<br><br>***YOUR TURN***<br>INPUT: {text}<br>OUTPUT: |
| **Long, basic** | ***INSTRUCTIONS***<br>You are an expert in the biomedical domain. You are a smart question answering assistant, specialized in answering biomedical questions. I will provide you the output format, the question to answer, and the abstract based on which to answer the question. |

| | |
|---|---|
| | Your answer must be one word: yes, no, or maybe. Do not provide any explanation or other characters.<br><br>***YOUR TURN***<br>INPUT: {text}<br>OUTPUT: |
| **Short, chat** | SYSTEM: You are an expert in the biomedical domain. You are a smart question answering assistant, specialized in answering biomedical questions. Answer based on the provided abstract. Only answer with yes, no, or maybe. Do not provide any explanation or other characters.<br><br>USER: {text} |
| **Long, chat** | SYSTEM: You are an expert in the biomedical domain. You are a smart question answering assistant, specialized in answering biomedical questions. I will provide you the output format, the question to answer, and the abstract based on which to answer the question.<br>Your answer must be one word: yes, no, or maybe. Do not provide any explanation or other characters.<br><br>USER: {text} |

#### RE – ChemProt

| Style | Template |
|---|---|
| **Short, basic** | ***INSTRUCTIONS***<br>You are an expert in the biomedical domain. You are a smart relation extraction assistant, specialized in relation extraction between a chemical entity and a gene entity. Classify relations between a chemical labeled as @CHEMICAL$ and a gene labeled as @GENE$. The answer must be one of CPR:3, CPR:4, CPR:5, CPR:6, CPR:9, or false. Do not provide any explanation or other characters.<br><br>***YOUR TURN***<br>INPUT: {text}<br>OUTPUT: |
| **Long, basic** | ***INSTRUCTIONS***<br>You are an expert in the biomedical domain. You are a smart relation extraction assistant, specialized in relation extraction between a chemical entity and a gene entity. Classify relations between a chemical labeled as @CHEMICAL$ and a gene labeled as @GENE$. I will provide you the output format, the definition of class categories, and the text based on which to answer.<br>Your answer must be one out of the six types of relations (CPR:3, CPR:4, CPR:5, CPR:6, CPR:9 or false) for the gene and chemical without any explanation or other characters.<br>Output Format:<br>Category Definition: |

| | |
|---|---|
| | CPR:3, which includes UPREGULATOR, ACTIVATOR, and INDIRECT UPREGULATOR<br>CPR:4, which includes DOWNREGULATOR, INHIBITOR ,and INDIRECT DOWNREGULATOR<br>CPR:5, which includes AGONIST, AGONIST ACTIVATOR, and AGONIST INHIBITOR<br>CPR:6, which includes ANTAGONIST<br>CPR:9, which includes SUBSTRATE, PRODUCT OF and SUBSTRATE PRODUCT OF<br>false, which indicates no relations<br><br>***YOUR TURN***<br>INPUT: {text}<br>OUTPUT: |
| **Short, chat** | SYSTEM: You are an expert in the biomedical domain. You are a smart relation extraction assistant, specialized in relation extraction between a chemical entity and a gene entity. Classify relations between a chemical labeled as @CHEMICAL$ and a gene labeled as @GENE$. The answer must be one of CPR:3, CPR:4, CPR:5, CPR:6, CPR:9, or false. Do not provide any explanation or other characters.<br><br>USER: {text} |
| **Long, chat** | SYSTEM: You are an expert in the biomedical domain. You are a smart relation extraction assistant, specialized in relation extraction between a chemical entity and a gene entity. Classify relations between a chemical labeled as @CHEMICAL$ and a gene labeled as @GENE$. I will provide you the output format, the definition of class categories, and the text based on which to answer.<br>Your answer must be one out of the six types of relations (CPR:3, CPR:4, CPR:5, CPR:6, CPR:9 or false) for the gene and chemical without any explanation or other characters.<br>Output Format:<br>Category Definition:<br>CPR:3, which includes UPREGULATOR, ACTIVATOR, and INDIRECT UPREGULATOR<br>CPR:4, which includes DOWNREGULATOR, INHIBITOR ,and INDIRECT DOWNREGULATOR<br>CPR:5, which includes AGONIST, AGONIST ACTIVATOR, and AGONIST INHIBITOR<br>CPR:6, which includes ANTAGONIST<br>CPR:9, which includes SUBSTRATE, PRODUCT OF and SUBSTRATE PRODUCT OF<br>false, which indicates no relations<br><br>USER: {text} |

#### RE – DDI

| Style | Template |
|---|---|
| **Short, basic** | ***INSTRUCTIONS***<br>You are an expert in the biomedical domain. You are a smart relation extraction assistant, specialized in relation extraction between drug entities. Classify relations between two drugs labeled as @DRUG$. The answer must be one of DDI-effect, DDI-mechanism, DDI-advise, DDI-false, or DDI-int. Do not provide any explanation or other characters.<br><br>***YOUR TURN***<br>INPUT: {text}<br>OUTPUT: |
| **Long, basic** | ***INSTRUCTIONS***<br>You are an expert in the biomedical domain. You are a smart relation extraction assistant, specialized in relation extraction between drug entities. Classify relations between two drugs labeled as @DRUG$. I will provide you the output format, the definition of class categories, and the text based on which to answer.<br>Your answer must be one out of the five types of relations (DDI-effect, DDI-mechanism, DDI-advise, DDI-false, and DDI-int) for the drugs without any explanation or other characters.<br>Output Format:<br>Category Definition:<br>DDI-mechanism: This type is used to annotate DDIs that are described by their PK mechanism (e.g. Grepafloxacin may inhibit the metabolism of theobromine)<br>DDI-effect: This type is used to annotate DDIs describing an effect (e.g. In uninfected volunteers, 46% developed rash while receiving SUSTIVA and clarithromycin) or a PD mechanism (e.g. Chlorthalidone may potentiate the action of other antihypertensive drugs)<br>DDI-advise: This type is used when a recommendation or advice regarding a drug interaction is given (e.g. UROXATRAL should not be used in combination with other alpha-blockers)<br>DDI-int: This type is used when a DDI appears in the text without providing any additional information (e.g. The interaction of omeprazole and ketoconazole has been established)<br>DDI-false, This type is used when no DDI relation appears<br><br>***YOUR TURN***<br>INPUT: {text}<br>OUTPUT: |
| **Short, chat** | SYSTEM: You are an expert in the biomedical domain. You are a smart relation extraction assistant, specialized in relation extraction between drug entities. Classify relations between two drugs labeled as @DRUG$. The answer must be one of DDI-effect, DDI-mechanism, |

| | |
|---|---|
| | DDI-advise, DDI-false, or DDI-int. Do not provide any explanation or other characters.<br><br>USER: {text} |
| **Long, chat** | SYSTEM: You are an expert in the biomedical domain. You are a smart relation extraction assistant, specialized in relation extraction between drug entities. Classify relations between two drugs labeled as @DRUG$. I will provide you the output format, the definition of class categories, and the text based on which to answer.<br>Your answer must be one out of the five types of relations (DDI-effect, DDI-mechanism, DDI-advise, DDI-false, and DDI-int) for the drugs without any explanation or other characters.<br>Output Format:<br>Category Definition:<br>DDI-mechanism: This type is used to annotate DDIs that are described by their PK mechanism (e.g. Grepafloxacin may inhibit the metabolism of theobromine)<br>DDI-effect: This type is used to annotate DDIs describing an effect (e.g. In uninfected volunteers, 46% developed rash while receiving SUSTIVA and clarithromycin) or a PD mechanism (e.g. Chlorthalidone may potentiate the action of other antihypertensive drugs)<br>DDI-advise: This type is used when a recommendation or advice regarding a drug interaction is given (e.g. UROXATRAL should not be used in combination with other alpha-blockers)<br>DDI-int: This type is used when a DDI appears in the text without providing any additional information (e.g. The interaction of omeprazole and ketoconazole has been established)<br>DDI-false, This type is used when no DDI relation appears<br><br>USER: {text} |

#### RE – GAD

| Style | Template |
|---|---|
| **Short, basic** | ***INSTRUCTIONS***<br>You are an expert in the biomedical domain. You are a smart relation extraction assistant, specialized in relation extraction between a disease entity and a gene entity. Classify relations between a disease labeled as @DISEASE$ and a gene labeled as @GENE$. Answer with 1 if there is a relation or 0 if there is not. Do not provide any explanation or other characters.<br><br>***YOUR TURN***<br>INPUT: {text}<br>OUTPUT: |
| **Long, basic** | ***INSTRUCTIONS***<br>You are an expert in the biomedical domain. You are a smart relation |

| | |
|---|---|
| | extraction assistant, specialized in relation extraction between a disease entity and a gene entity. Classify relations between a disease labeled as @DISEASE$ and a gene labeled as @GENE$. I will provide you the output format, the definition of class categories, and the text based on which to answer.<br>Answer with 1 or 0. Do not provide any explanation or other characters.<br>Output Format:<br>Category Definition:<br>0: there is no relation between the disease entity and the gene entity<br>1: there is a relation between the disease entity and the gene entity<br><br>***YOUR TURN***<br>INPUT: {text}<br>OUTPUT: |
| **Short, chat** | SYSTEM: You are an expert in the biomedical domain. You are a smart relation extraction assistant, specialized in relation extraction between a disease entity and a gene entity. Classify relations between a disease labeled as @DISEASE$ and a gene labeled as @GENE$. Answer with 1 if there is a relation or 0 if there is not. Do not provide any explanation or other characters.<br><br>USER: {text} |
| **Long, chat** | SYSTEM: You are an expert in the biomedical domain. You are a smart relation extraction assistant, specialized in relation extraction between a disease entity and a gene entity. Classify relations between a disease labeled as @DISEASE$ and a gene labeled as @GENE$. I will provide you the output format, the definition of class categories, and the text based on which to answer.<br>Answer with 1 or 0. Do not provide any explanation or other characters.<br>Output Format:<br>Category Definition:<br>0: there is no relation between the disease entity and the gene entity<br>1: there is a relation between the disease entity and the gene entity<br><br>USER: {text} |

#### SS – BIOSSES

| Style | Template |
|---|---|
| **Short, basic** | ***INSTRUCTIONS***<br>You are an expert in the biomedical domain. You are a smart semantic similarity scoring assistant, specialized in the scoring of semantic similarity between two sentences. Score the semantic similarity between two sentences named sentence1 and sentence2. Your score must be a number from 0.0 to 4.0 signifying the similarity between two sentences. Do not provide any explanation or other characters. |

| | |
|---|---|
| | ***YOUR TURN***<br>INPUT: {text}<br>OUTPUT: |
| **Long, basic** | ***INSTRUCTIONS***<br>You are an expert in the biomedical domain. You are a smart semantic similarity scoring assistant, specialized in the scoring of semantic similarity between two sentences. Score the semantic similarity between two sentences named sentence1 and sentence2. I will provide you the output format, the definition of the increments on the scoring scale, and the two sentences based on which to answer. Your answer is the semantic similarity expressed as a continuous number from 0.0 (no relation) to 4.0 (equivalent) without any explanation or other characters.<br>Output Format:<br>Category Definition:<br>0.0, the two sentences are on different topics.<br>1.0, the two sentences are not equivalent, but are on the same topic.<br>2.0, the two sentences are not equivalent, but share some details.<br>3.0, the two sentences are roughly equivalent, but some important information differs/missing.<br>4.0, The two sentences are completely or mostly equivalent, as they mean the same thing. Note that fractions (e.g. 1.7 or 3.5) are also valid answers.<br><br>***YOUR TURN***<br>INPUT: {text}<br>OUTPUT: |
| **Short, chat** | SYSTEM: You are an expert in the biomedical domain. You are a smart semantic similarity scoring assistant, specialized in the scoring of semantic similarity between two sentences. Score the semantic similarity between two sentences named sentence1 and sentence2. Your score must be a number from 0.0 to 4.0 signifying the similarity between two sentences. Do not provide any explanation or other characters.<br><br>USER: {text} |
| **Long, chat** | SYSTEM: You are an expert in the biomedical domain. You are a smart semantic similarity scoring assistant, specialized in the scoring of semantic similarity between two sentences. Score the semantic similarity between two sentences named sentence1 and sentence2. I will provide you the output format, the definition of the increments on the scoring scale, and the two sentences based on which to answer. Your answer is the semantic similarity expressed as a continuous number from 0.0 (no relation) to 4.0 (equivalent) without any explanation or other characters.<br>Output Format:<br>Category Definition: |

| | |
|---|---|
| | 0.0, the two sentences are on different topics.<br>1.0, the two sentences are not equivalent, but are on the same topic.<br>2.0, the two sentences are not equivalent, but share some details.<br>3.0, the two sentences are roughly equivalent, but some important information differs/missing.<br>4.0, The two sentences are completely or mostly equivalent, as they mean the same thing. Note that fractions (e.g. 1.7 or 3.5) are also valid answers.<br><br>USER: {text} |

#### Verbal Confidence Elicitation

Prompt templates are also required when verbally eliciting confidence scores from LLMs (see Appendix 2). Here too we need *basic*- and *chat-targeted* templates, depending on the type of LLM:

| Style | Template |
|---|---|
| **Basic** | $\{p\}$<br>OUTPUT: $\{y\}$<br>How confident are you of your previous answer? Reply with only a float between 0.0 and 1.0, where 0.0 means you are completely unsure of your answer and 1.0 means you are completely sure. |
| **Chat** | SYSTEM: $\{p_{instructions}\}$<br>USER: $\{p_{text}\}$<br>ASSISTANT: $\{y\}$<br>USER: How confident are you of your previous answer? Reply with only a float between 0.0 and 1.0, where 0.0 means you are completely unsure of your answer and 1.0 means you are completely sure. |

where $p$ is the unique prompt submitted to the LLM, $y$ is the response, $p_{instructions}$ is the instructions portion of $p$, which is constant for each dataset, and $p_{text}$ is the text of each example in the dataset, which makes $p$ unique.
