## Supplemental File 2 for "A Study of Calibration as a Measurement of Trustworthiness of Large Language Models in Biomedical Research"

### Supplementary File 2 – Formulae

This is a supplementary file to a full manuscript *A Study of Calibration as a Measurement of Trustworthiness of Large Language Models in Biomedical Research*.

---

#### Similarity Functions

The below similarity functions were used during confidence assignment as well as during evaluation. They all return a degree of similarity in [0, 1].

##### Equality

Defined as:

$$eq(a, b) = \begin{cases} 1 & if\ a = b \\ 0 & if\ a \neq b \end{cases}$$

where $a$ and $b$ are objects of any type.

##### Jaccard

Defined as:

$$j(A, B) = \frac{|A \cap B|}{|A \cup B|}$$

where A and B are sets.

##### F1 (macro)

Defined as:

$$\mathrm{f1}(A, B) = \frac{2 \times TP}{2 \times TP + FP + FN}$$

where A and B are lists, and TP/FP/FN are the element-wise exact matches between elements of the lists.

##### Relative Distance

Defined as:

$$rdist(a, b) = 1 - \frac{|a - b|}{a + b}$$

where $a$ and $b$ are numerical values.

### Confidence Score Assignment

Below we define the three methods we employed in this study to assign confidence scores to LLM responses.

#### Verbal

We define verbal confidence score $c_v$ as:

$$c_v = p_v(p, y)$$

where $p_v$ is a prompt to verbally elicit a confidence score from the LLM given the original prompt $p$ and the response $y$ to $p$. Prompt details in Appendix 1.

#### Self-consistency

We define the self-consistency confidence score $c_{sc}$ as:

$$c_{sc} = \frac{1}{M}\sum_{i=1}^{M} s(y, \tilde{Y}_i)$$

where $y$ is the main response, $\tilde{Y}$ is the set of $M$ candidate responses to be compared with $y$ using the appropriate similarity function $s$.

#### Hybrid

We define the hybrid confidence score $c_h$ as:

$$c_h = \frac{1}{M}\sum_{i=1}^{M} \begin{cases} (c_v + \tilde{C}_i)/\,2 & if\ s(y, \tilde{Y}_i) \geq \mathrm{T} \\ 1 - c_v & if\ s(y, \tilde{Y}_i) < \mathrm{T} \end{cases}$$

where $c_v$ is the verbal confidence score, $\tilde{C}$ is the set of $M$ candidate verbal confidence scores, $y$ is the main response, $\tilde{Y}$ is the set of $M$ candidate responses, $sim$ the appropriate similarity function and $\mathrm{T}$ the similarity threshold, where $\mathrm{T} \in [0, 1]$.

### Metrics

The below metric was used to evaluate LLMs.

#### Flexible Expected Calibration Error (Flex-ECE)

Defined as:

$$ECE_{Flex} = \sum_{m=1}^{M} \frac{|B_m|}{n} |sim(B_m) - conf(B_m)|$$

where $B$ is a bin of $< response, target, confidence\ score >$ tuples, $M$ is the number of bins, $n$ the size of the dataset, $sim$ is the average similarity in the bin, and $conf$ the average confidence in the bin. More formally, $sim$ is defined as:

$$sim(B) = \frac{1}{|B_m|} \sum_{i \in B_m} s(y_i, \tilde{y}_i)$$

where $y$ is a response and $\tilde{y}$ is the target response. And $conf$ is defined as:

$$conf(B) = \frac{1}{|B_m|} \sum_{i \in B_m} c_i$$

where $c$ is a confidence score.
