## Supplement File 3 for "A Study of Calibration as a Measurement of Trustworthiness of Large Language Models in Biomedical Research"

### Supplementary File 3 – Additional Tables and Figures

This is a supplementary file to a full manuscript *A Study of Calibration as a Measurement of Trustworthiness of Large Language Models in Biomedical Research*.

---

#### Datasets

**Supplementary Table S1.** A description of the format of responses in each dataset, as well as the similarity function used to compare responses. Similarity functions are described in Supplementary File 2.

| Dataset | Format | Connotation | Similarity Function |
|---|---|---|---|
| **BC2GM** | Set of substrings of input string | Entities found in the text | Jaccard |
| **BC5-disease** | | | |
| **NCBI Disease** | | | |
| **JNLPBA** | | | |
| **BC5-chem** | | | |
| **EBM-PICO** | Array of same length as tokens in text, where each element is one of {I-PAR, I-INT, I-OUT, O}, for participant, intervention, outcome and none, respectively | | F1 (macro) |
| **ChemProt** | One string from a pre-defined set of strings | Relationship between (pre-annotated) entities in the text | Equality |
| **DDI** | | | |
| **GAD** | | | |
| **BIOSSES** | Float between 0.0 and 4.0 | The degree of similarity between sentences | Relative distance |
| **HoC** | Subset of pre-defined set of strings | Pre-defined set of hallmarks of cancer | Jaccard |
| **BioASQ** | Yes/No | The answer to the question | Equality |
| **PubMedQA** | Yes/No/Maybe | | |

#### Results

This section contains diagrams of results, before and after calibration.

In the tables and figures below, the following abbreviations are used.

- Tasks:
  - DC: Document classification
  - NER: Named-entity recognition
  - QA: Question answering
  - PICO: Population, Intervention, Comparator, Outcome
  - SS: Sentence similarity
- Calibration Methods:
  - HB: Histogram binning
  - IR: Isotonic regression
- Models:
  - F-XXL: Flan-T5-XXL
  - G-3: GPT-3.5-Turbo
  - G-4: GPT-4
  - L-8: Meta-Llama-3-8B-Instruct
  - ML-13: MedLlama-13B
  - ML-8: Medicine-Llama3-8B
  - Y-34: Yi-1.5-34B-Chat
  - Z-7: Zephyr-7B-Beta

### Before Calibration

#### Reliability Diagrams

Below is a set of reliability diagrams for each dataset. Each set contains sub-diagrams for each model and strategy. Reliability diagrams are visualizations of how calibrated a model is, by plotting the bin-wise accuracies (blue bars) and accuracy gaps (red bars) to the ideal bin-wise accuracy (the diagonal dotted line).

In the following diagrams, "Success" refers to the percentage of examples for which a valid confidence score was returned by the model. Since LLMs were not forced to generate valid verbal confidence scores, responses that could not be automatically parsed into a numerical value were labelled *unsuccessful* and were ignored. We arbitrarily establish a 50% success rate threshold to calculate Flex-ECE and plot reliability diagrams to ensure they are comparable across models and datasets.

#### *DC – HoC*

**Supplementary Figure S1**. Reliability diagrams for all LLMs and confidence assignment strategies on the HoC (Hallmarks of Cancer) dataset.

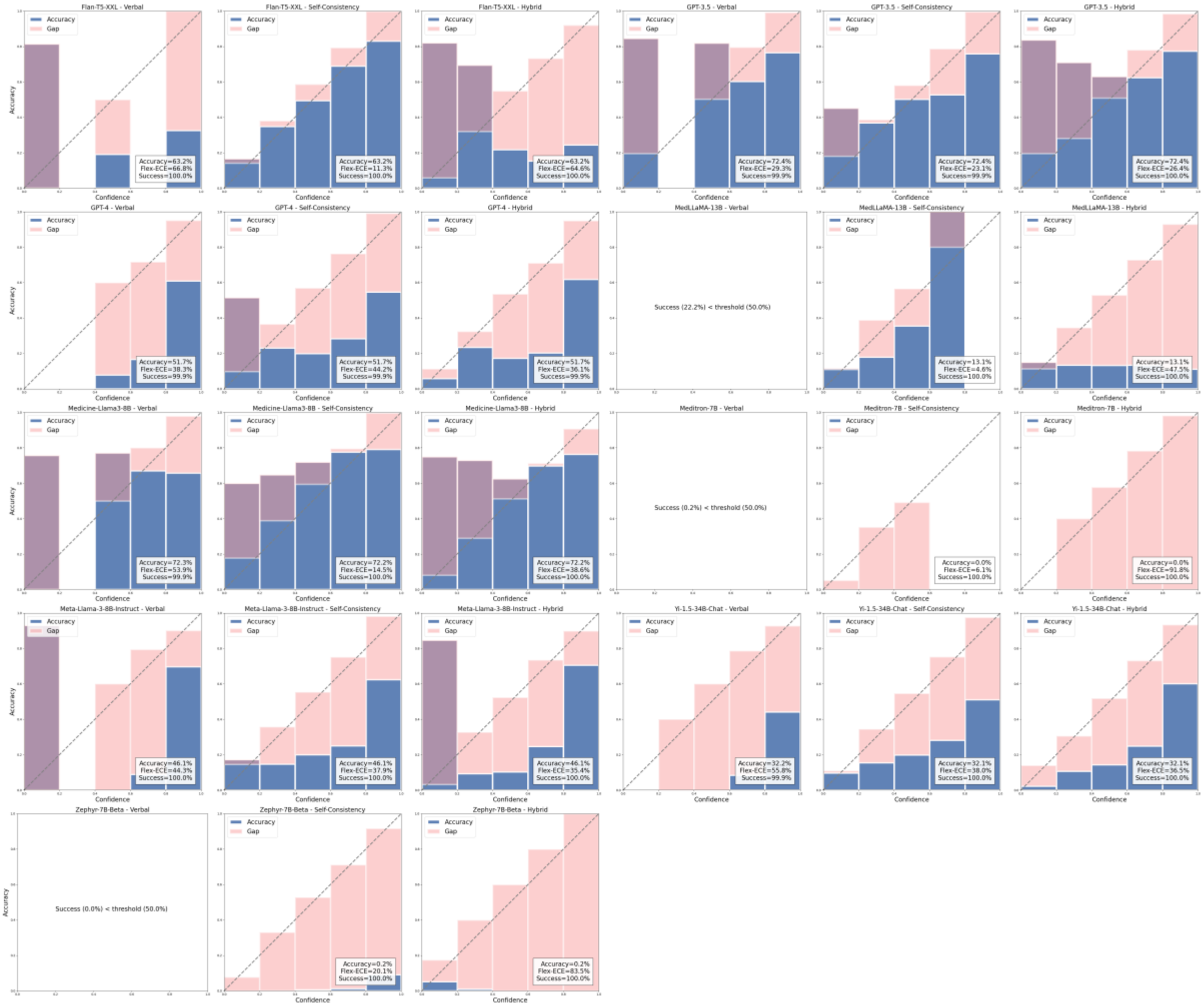

#### *NER – BC2GM*

**Supplementary Figure S2**. Reliability diagrams for all LLMs and confidence assignment strategies on the BC2GM dataset.

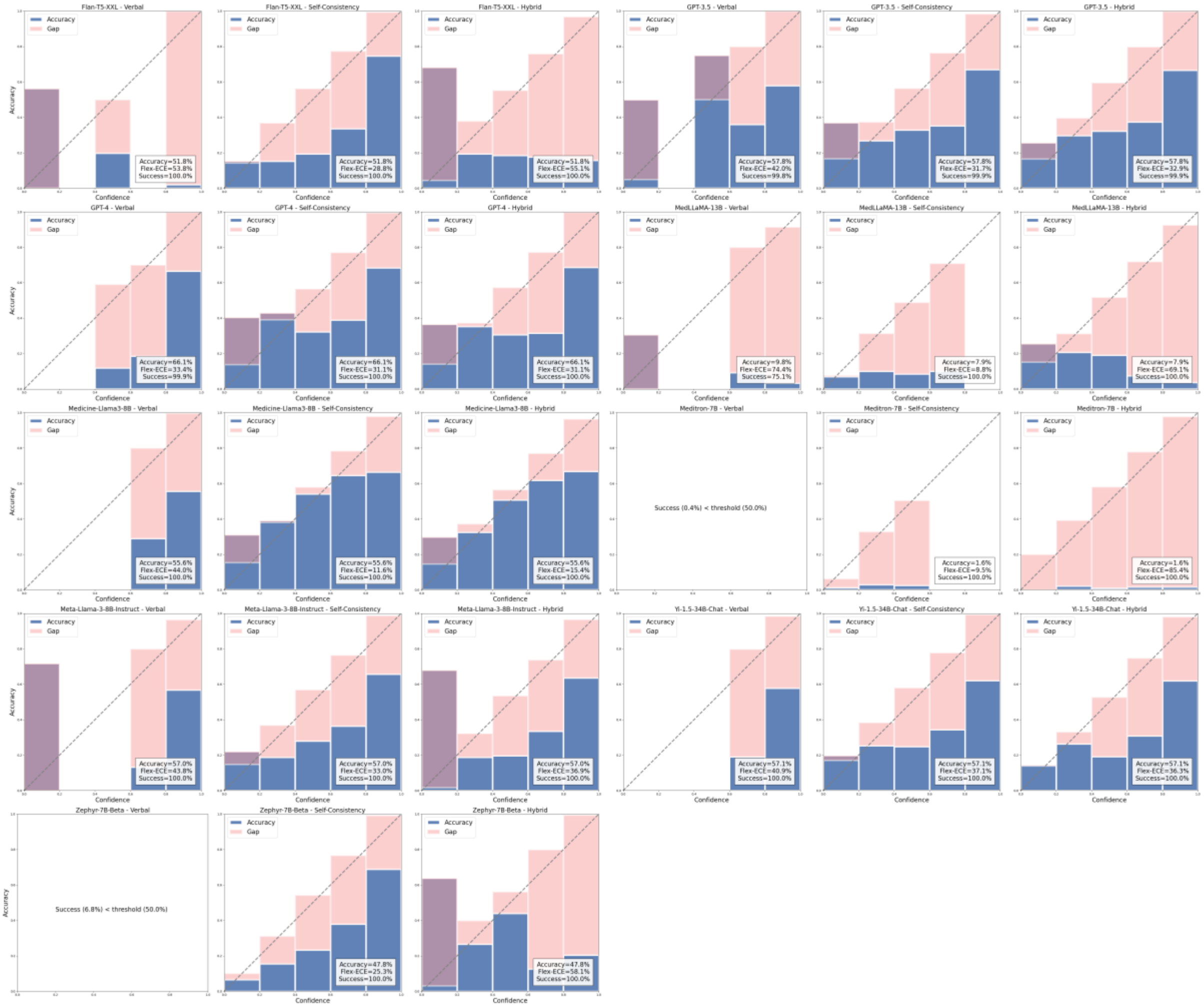

#### *NER – BC5-Chemical*

**Supplementary Figure S3**. Reliability diagrams for all LLMs and confidence assignment strategies on the BC5-Chemical dataset.

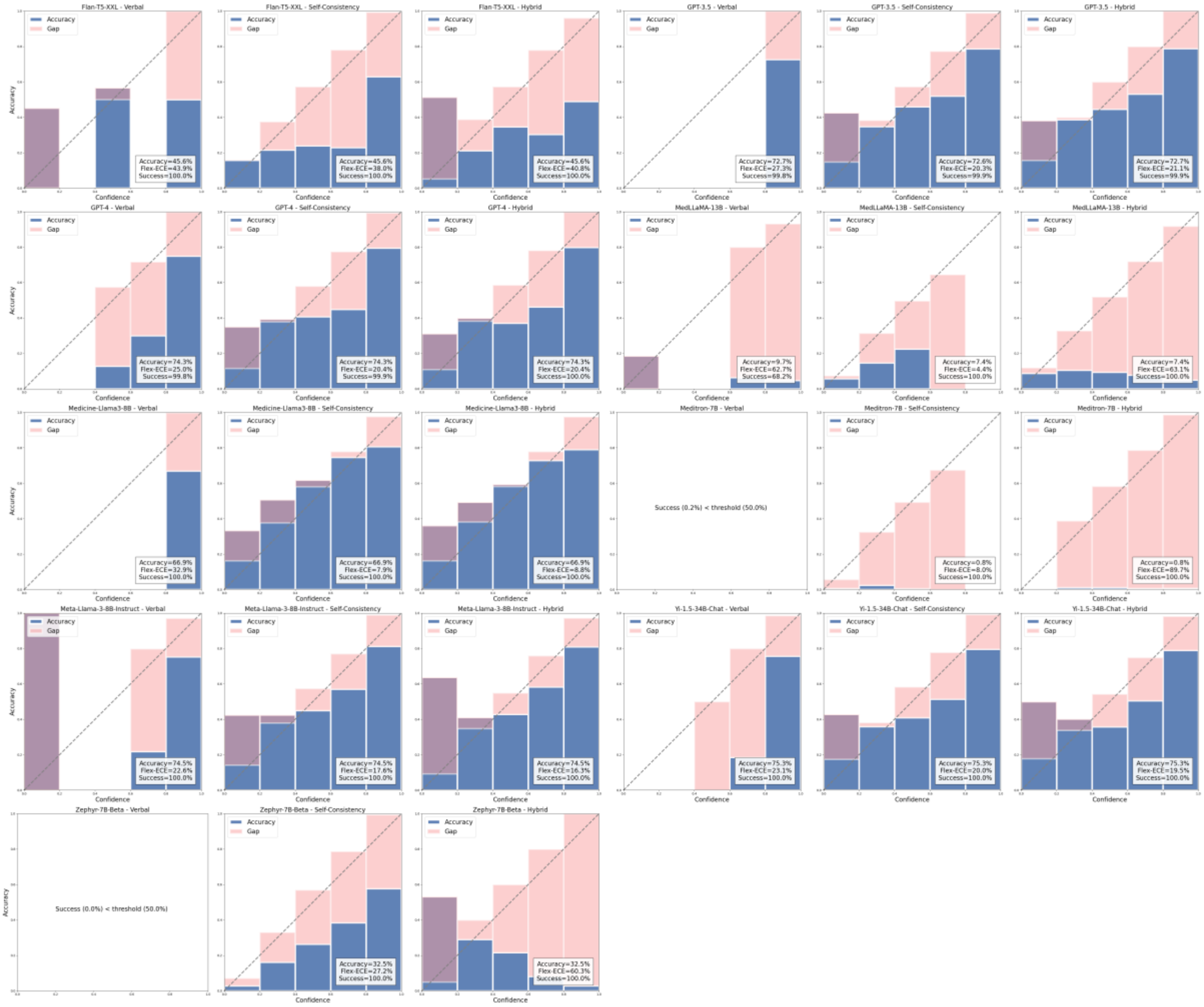

#### *NER – BC5-Disease*

**Supplementary Figure S4**. Reliability diagrams for all LLMs and confidence assignment strategies on the BC5-Disease dataset.

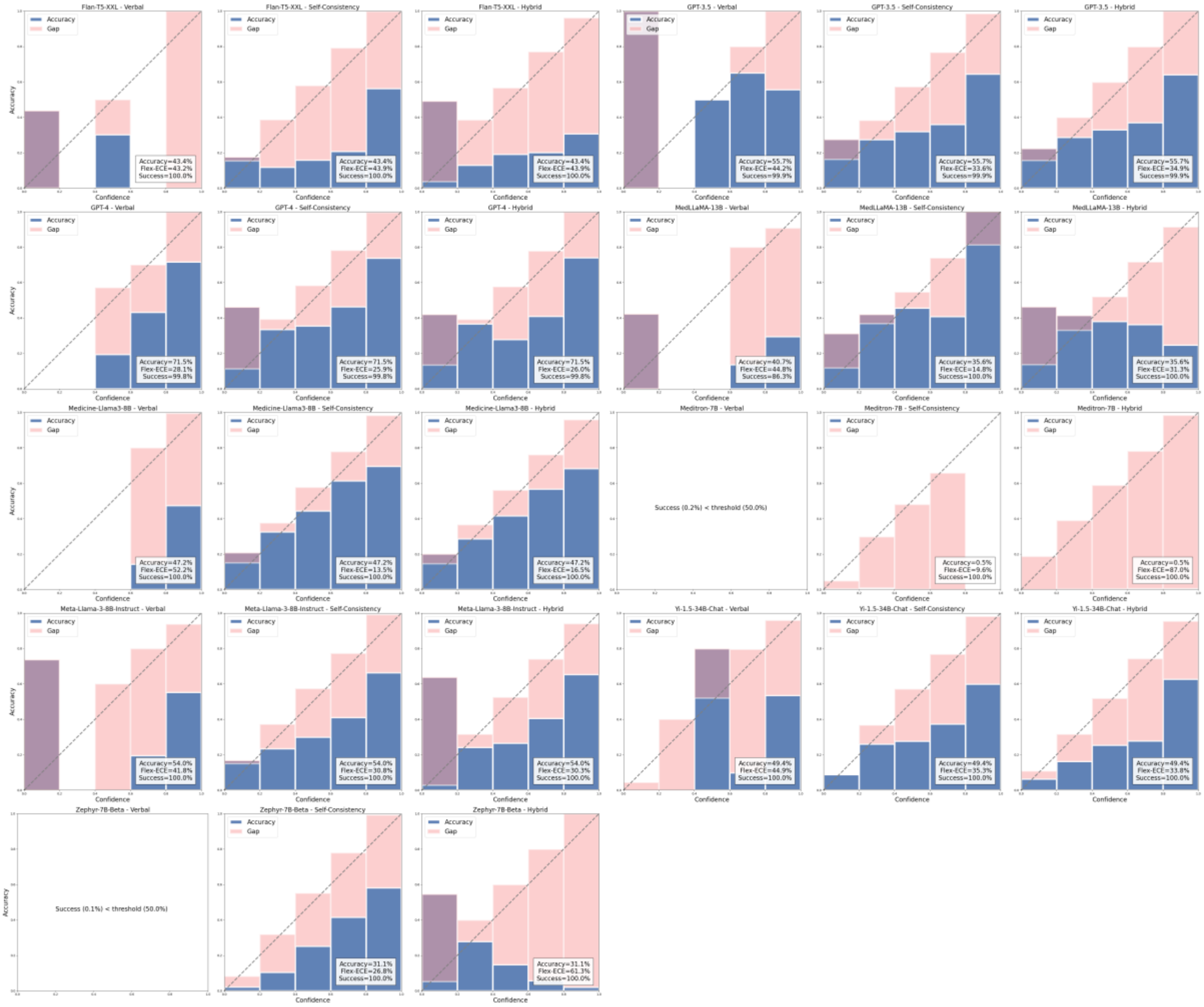

*NER – JNLPBA*

**Supplementary Figure S5**. Reliability diagrams for all LLMs and confidence assignment strategies on the JNLPBA dataset.

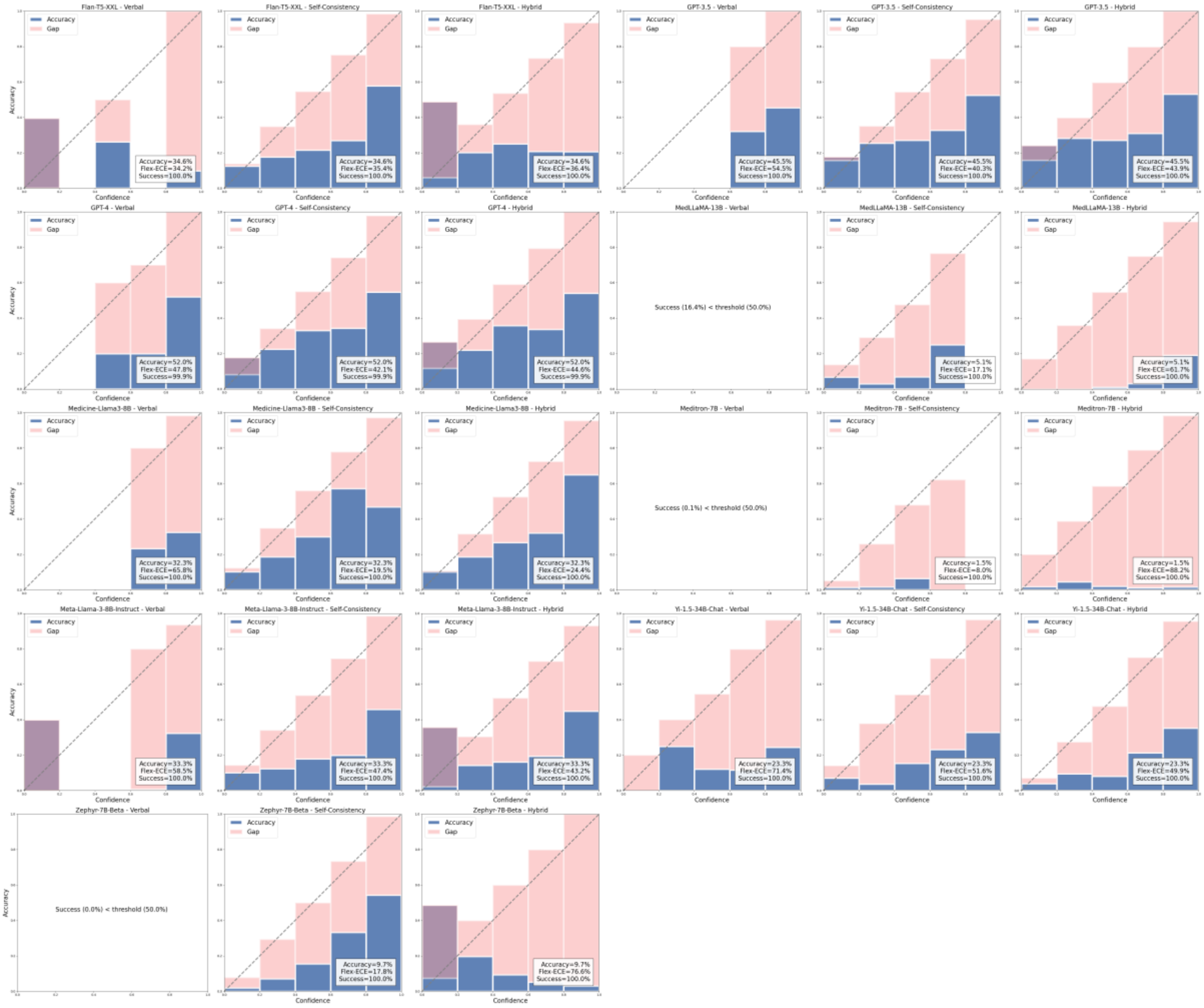

#### *NER – NCBI-Disease*

**Supplementary Figure S6.** Reliability diagrams for all LLMs and confidence assignment strategies on the NCBI-Disease dataset.

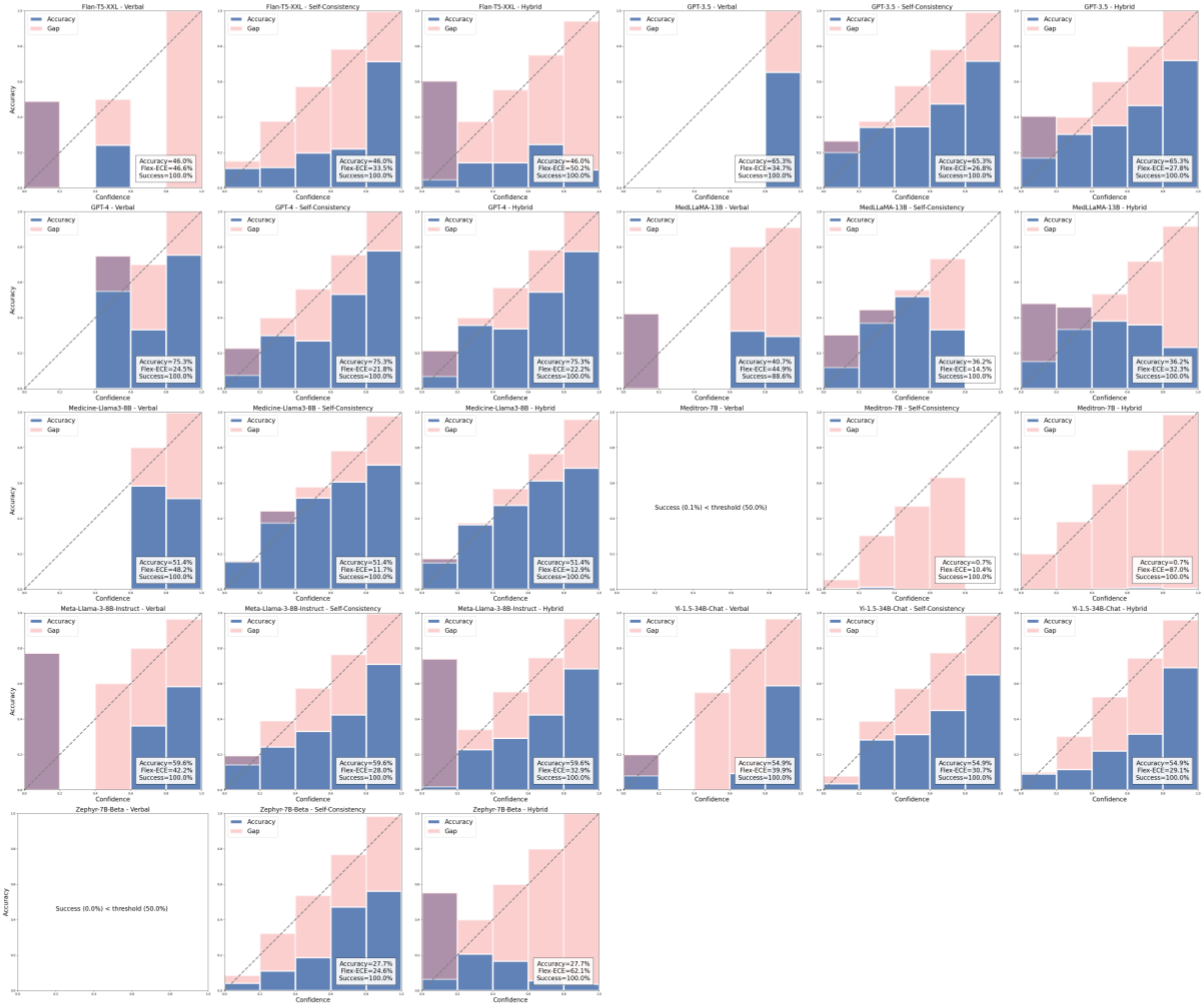

#### *PICO – EBM-PICO*

**Supplementary Figure S7**. Reliability diagrams for all LLMs and confidence assignment strategies on the EBM-PICO dataset.

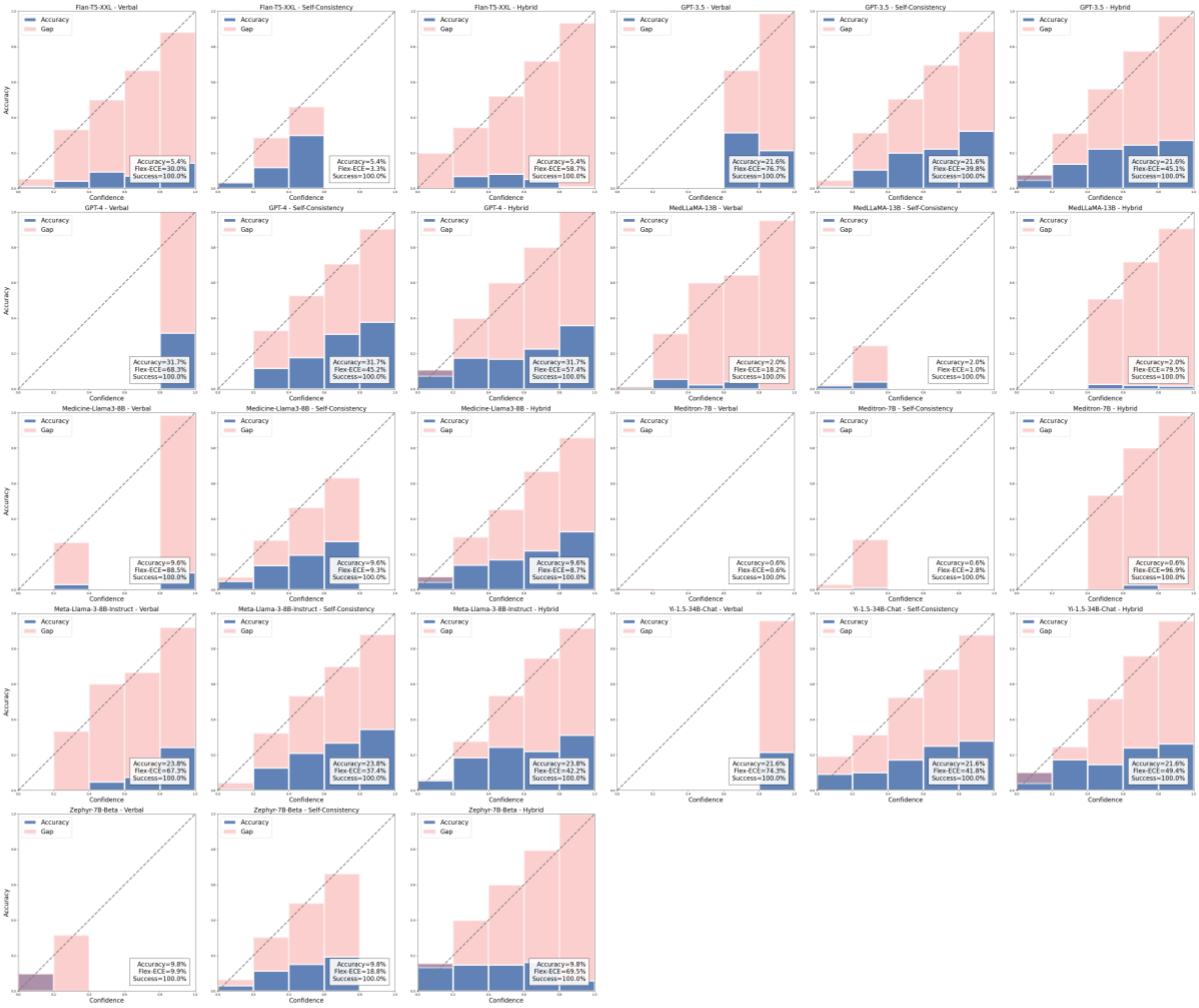

#### *QA – BioASQ*

**Supplementary Figure S8.** Reliability diagrams for all LLMs and confidence assignment strategies on the BioASQ dataset.

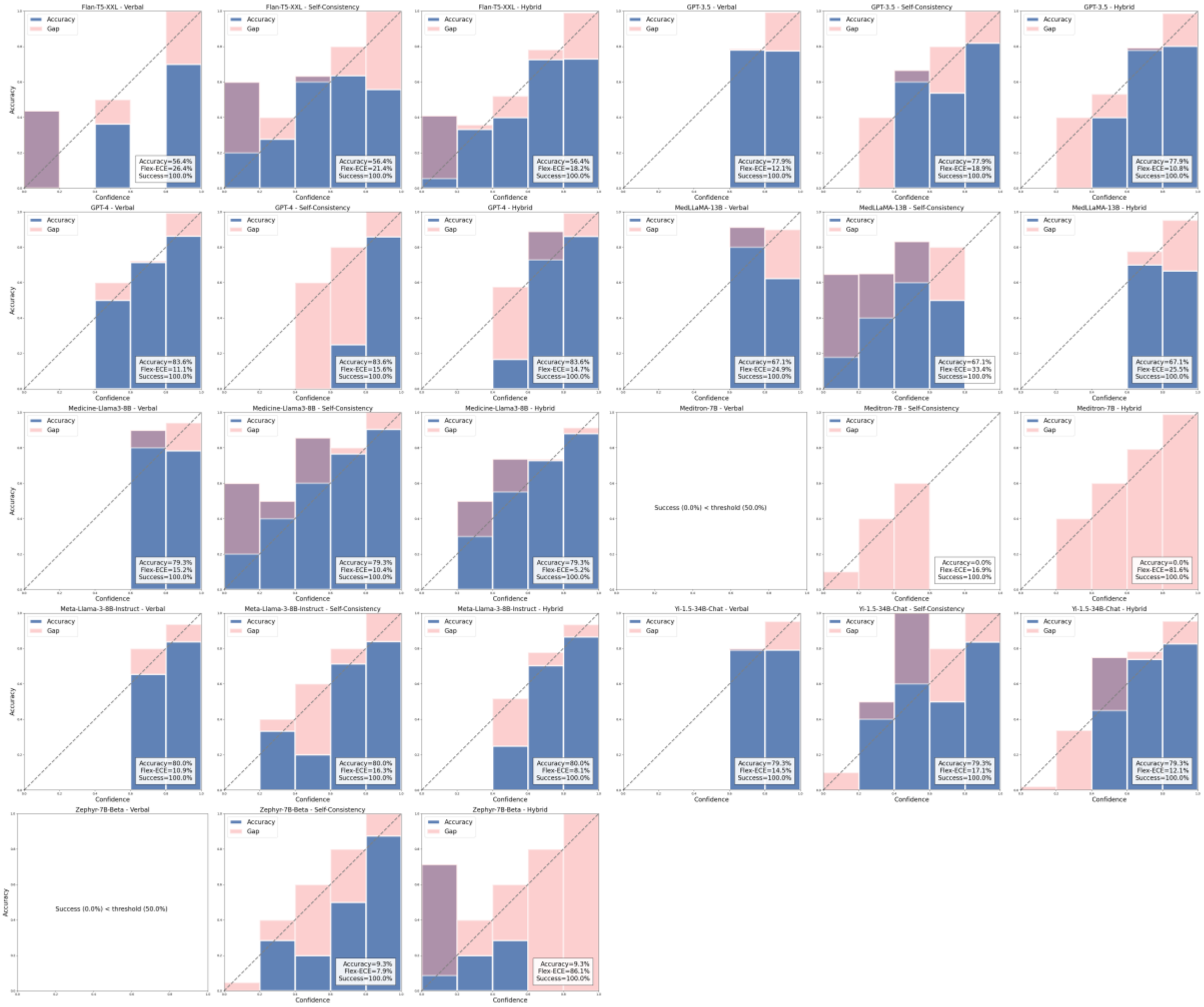

#### QA – PubMedQA

**Supplementary Figure S9**. Reliability diagrams for all LLMs and confidence assignment strategies on the PubMedQA dataset.

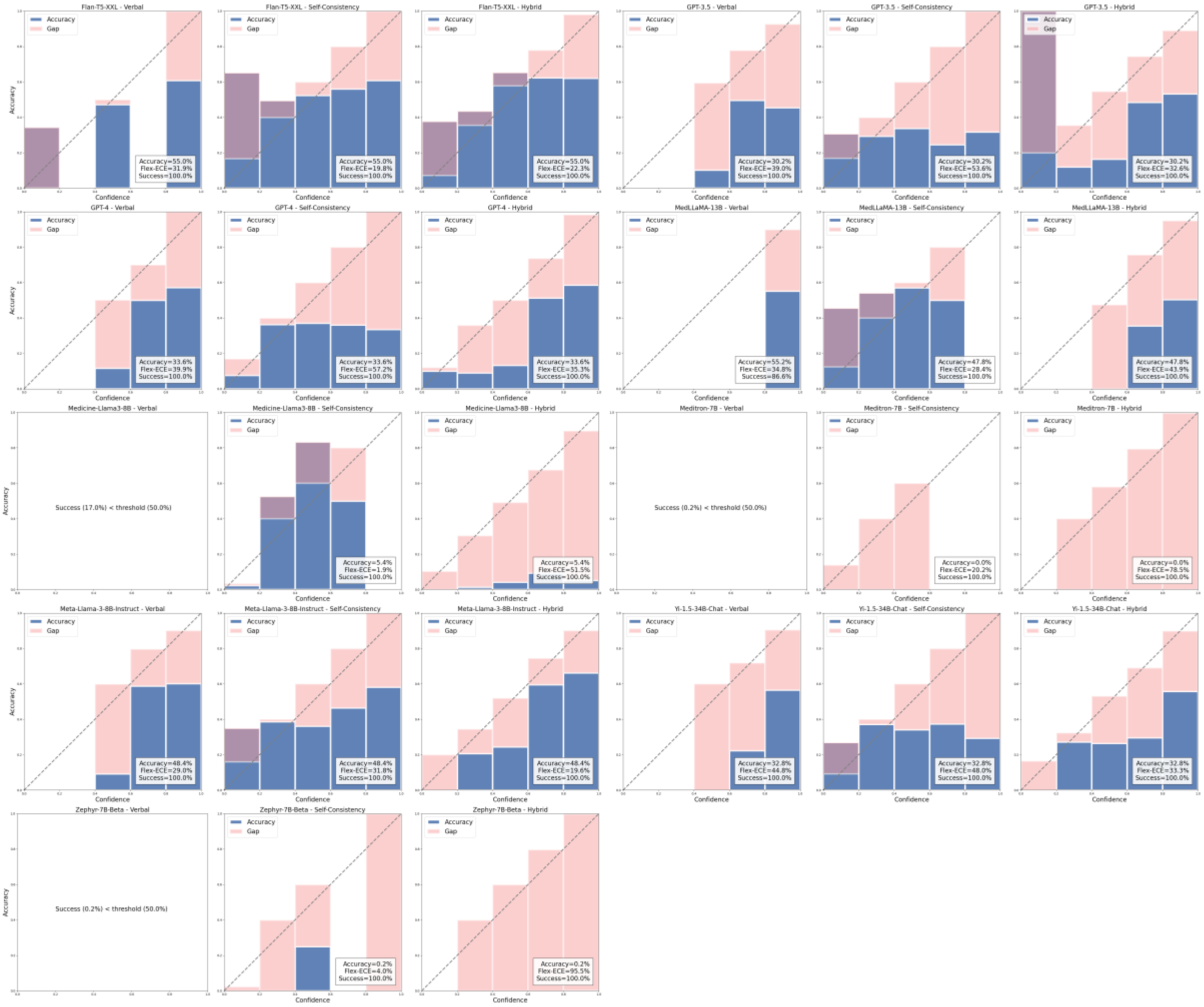

#### *RE – ChemProt*

**Supplementary Figure S10**. Reliability diagrams for all LLMs and confidence assignment strategies on the ChemProt dataset.

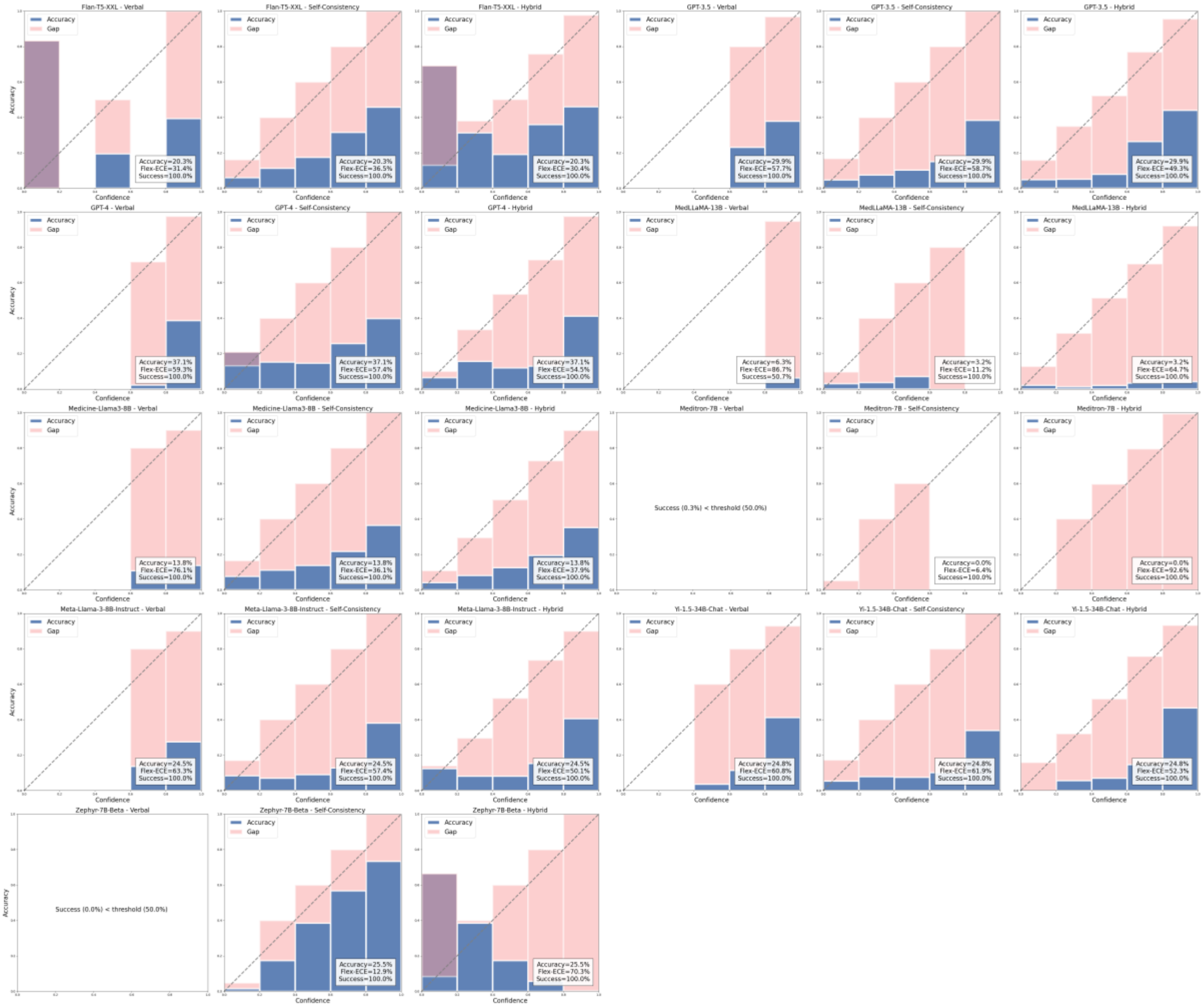

*RE – DDI*

**Supplementary Figure S11.** Reliability diagrams for all LLMs and confidence assignment strategies on the DDI dataset.

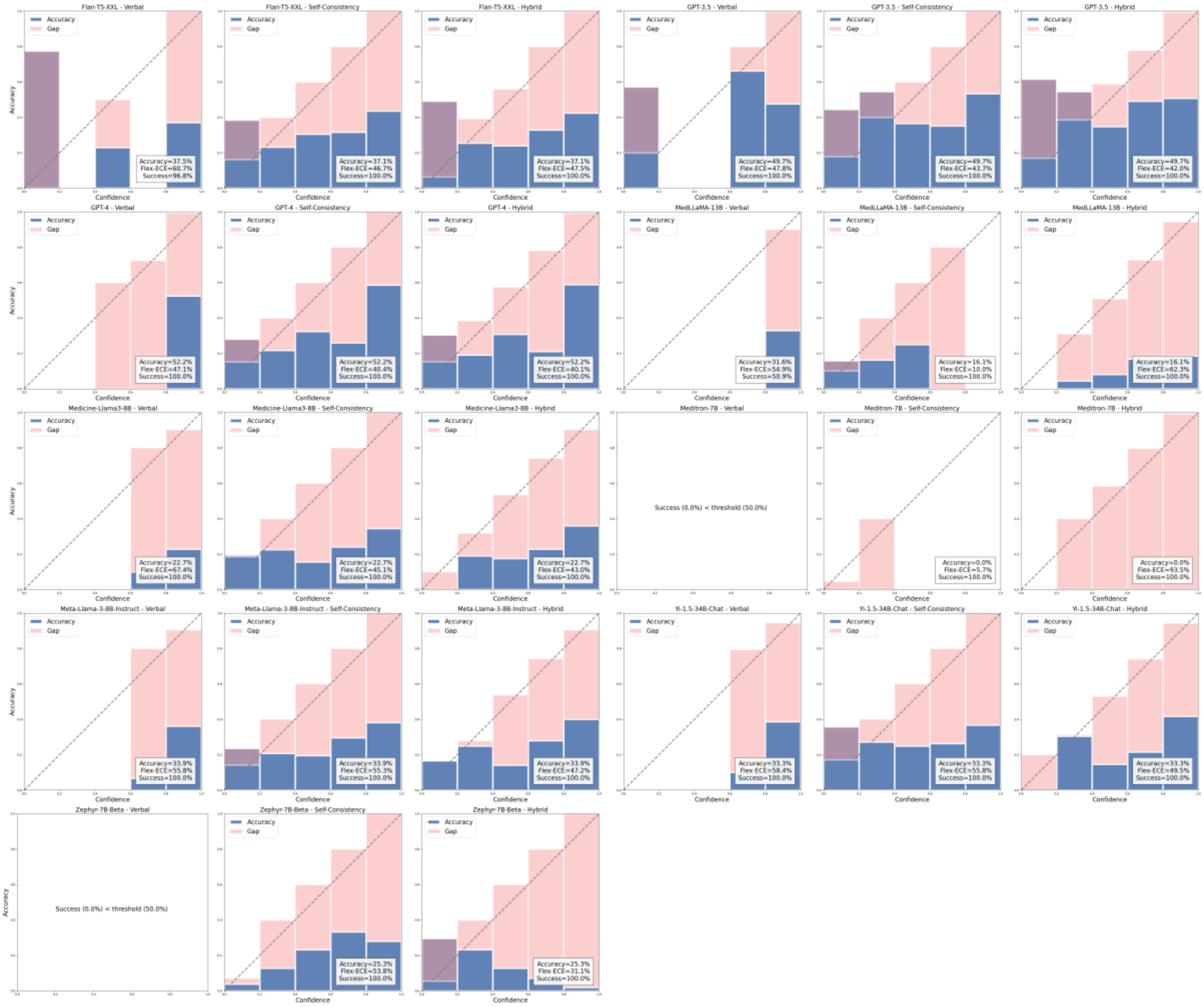

#### *RE – GAD*

**Supplementary Figure S12**. Reliability diagrams for all LLMs and confidence assignment strategies on the GAD dataset.

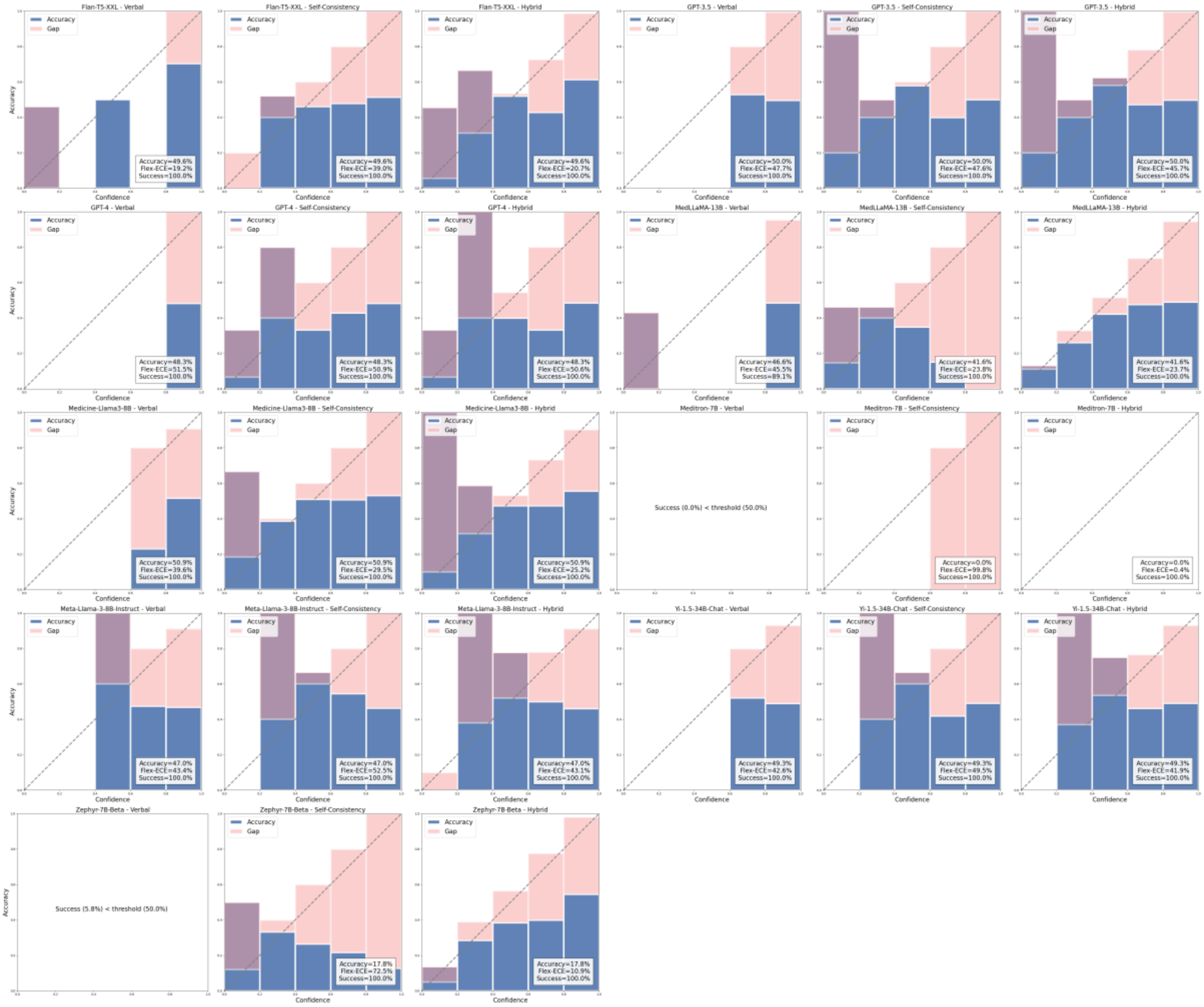

*SS – BIOSSES*

**Supplementary Figure S13**. Reliability diagrams for all LLMs and confidence assignment strategies on the BIOSSES dataset.

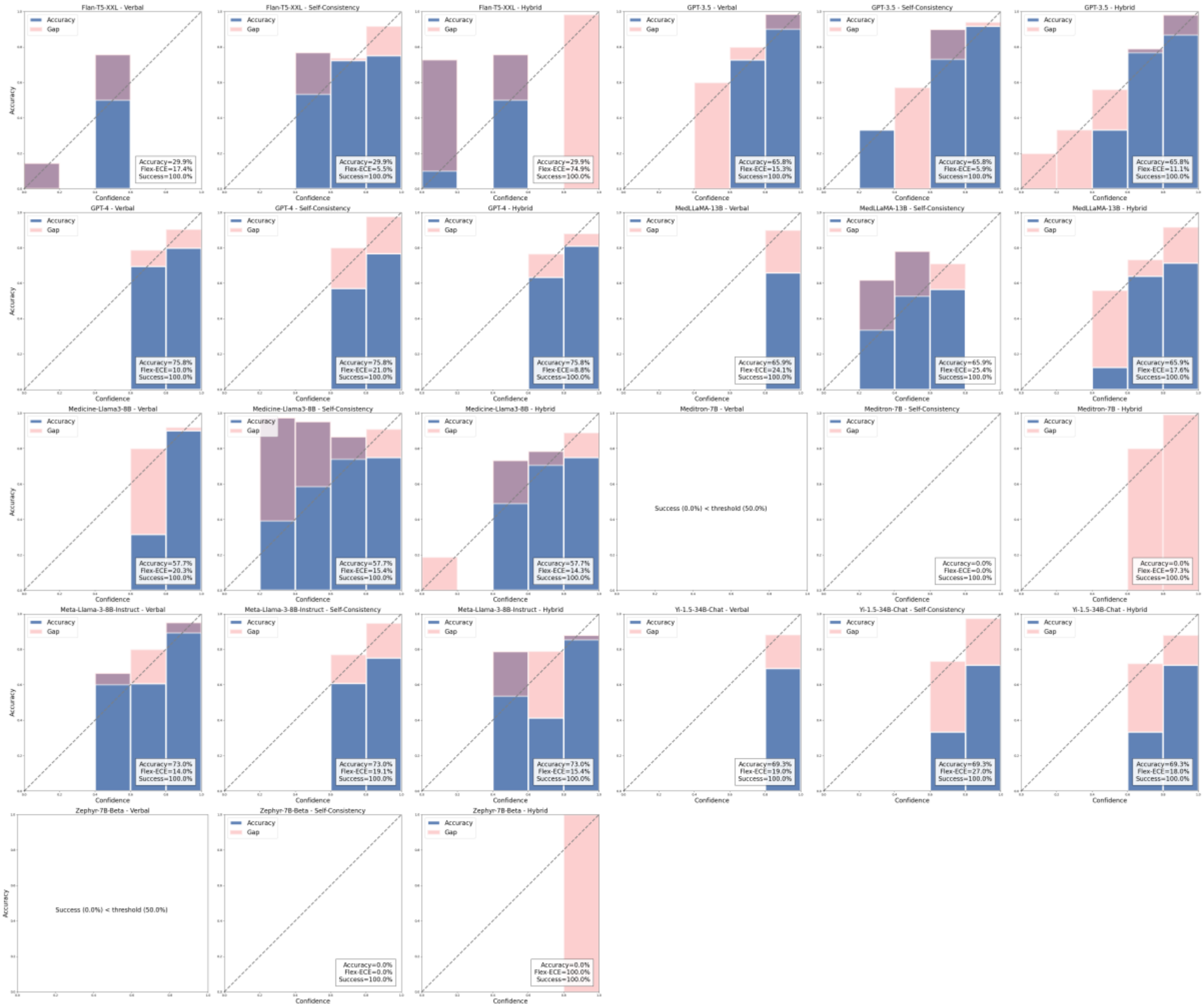

#### Flex ECE

**Supplementary Figure S14**. The full set of Flex-ECE scores, arranged by LLM and strategy. Horizontal dotted lines denote the median Flex-ECE score for the entire dataset.

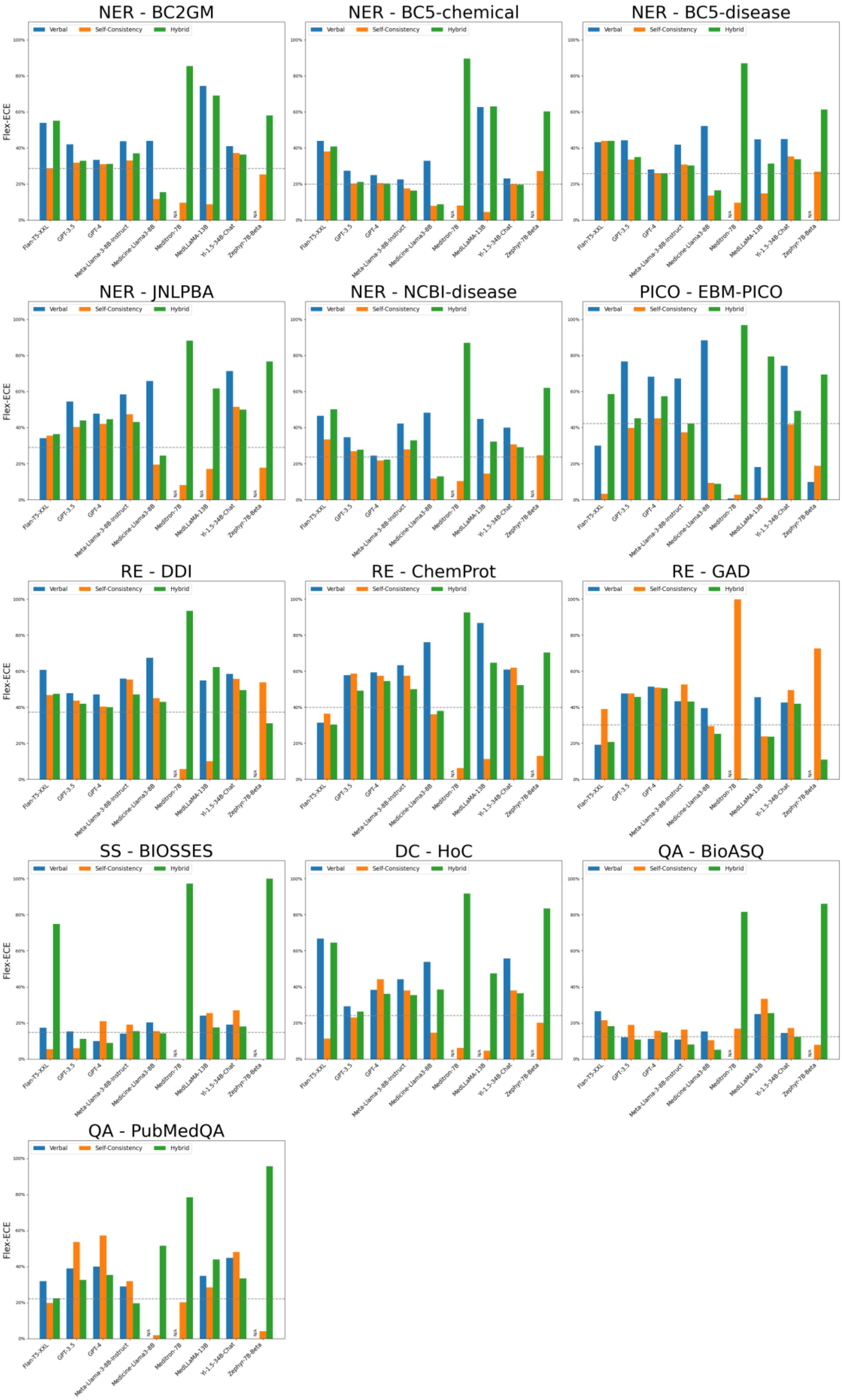
NER - BC2GM
NER - BC5-chemical
NER - BC5-disease
NER - JNLPBA
NER - NCBI-disease
PICO - EBM-PICO
RE - DDI
RE - ChemProt
RE - GAD
SS - BIOSSES
DC - HoC
QA - BioASQ
QA - PubMedQA
Verbal
Self-Consistency
Hybrid
Flex-ECE
0%
20%
40%
60%
80%
100%
Flan-T5-XXL
GPT-3.5
GPT-4
Meta-Llama-3-8B-Instruct
Medicine-Llama3-8B
Meditron-7B
MedLLaMA-13B
Yi-1.5-34B-Chat
Zephyr-7B-Beta
N/A

**Supplementary Table S2**. Flex-ECE scores by dataset and model using the verbal confidence assignment strategy. Only task-model-strategy combinations with a success rate ≥50% are included.

| Task | Dataset | F-XXL | G-3 | G-4 | ML-13 | ML-8 | M-7 | L-8 | Y-34 | Z-7 | Mean |
|---|---|---|---|---|---|---|---|---|---|---|---|
| DC | HoC | 66.8 | 29.3 | 38.3 | - | 53.9 | - | 44.3 | 55.8 | - | 48.1 |
| | BC2GM | 53.8 | 42.1 | 33.4 | 74.4 | 44.0 | - | 43.8 | 40.9 | - | 47.5 |
| | BC5-chemical | 43.9 | 27.3 | 25.0 | 62.7 | 32.9 | - | 22.6 | 23.1 | - | 33.9 |
| | BC5-disease | 43.2 | 44.2 | 28.1 | 44.8 | 52.2 | - | 41.8 | 44.9 | - | 42.7 |
| | JNLPBA | 34.2 | 54.5 | 47.8 | - | 65.8 | - | 58.5 | 71.4 | - | 55.4 |
| NER | NCBI-disease | 46.6 | 34.7 | 24.5 | 44.9 | 48.2 | - | 42.2 | 39.9 | - | 40.1 |
| PICO | EBM-PICO | 30.0 | 76.7 | 68.3 | 18.2 | 88.5 | 0.6 | 67.3 | 74.3 | 9.9 | 48.2 |
| | BioASQ | 26.4 | 12.1 | 11.1 | 24.9 | 15.2 | - | 10.9 | 14.5 | - | 16.4 |
| QA | PubMedQA | 31.9 | 39.0 | 39.9 | 34.8 | - | - | 29.0 | 44.8 | - | 36.6 |
| | ChemProt | 31.4 | 57.7 | 59.3 | 86.7 | 76.1 | - | 63.3 | 60.8 | - | 62.2 |
| | DDI | 60.7 | 47.8 | 47.1 | 54.9 | 67.4 | - | 55.9 | 58.4 | - | 56.0 |
| RE | GAD | 19.2 | 47.7 | 51.4 | 45.5 | 39.6 | - | 43.4 | 42.6 | - | 41.3 |
| SS | BIOSSES | 17.4 | 15.3 | 10.0 | 24.1 | 20.3 | - | 14.0 | 19.0 | - | 17.1 |
| | **Mean** | 38.9 | 40.6 | 37.2 | 46.9 | 50.3 | 0.6 | 41.3 | 45.4 | 9.9 | 42.0 |

**Supplementary Table S3**. Flex-ECE scores by dataset and model using the self-consistency confidence assignment strategy. Only task-model-strategy combinations with a success rate ≥50% are included.

| Task | Dataset | F-XXL | G-3 | G-4 | ML-13 | ML-8 | M-7 | L-8 | Y-34 | Z-7 | Mean |
|---|---|---|---|---|---|---|---|---|---|---|---|
| DC | HoC | 11.3 | 23.1 | 44.2 | 4.6 | 14.5 | 6.1 | 37.9 | 38.0 | 20.1 | 22.2 |
| | BC2GM | 28.8 | 31.8 | 31.1 | 8.8 | 11.6 | 9.5 | 33.0 | 37.1 | 25.3 | 24.1 |
| | BC5-chemical | 38.0 | 20.3 | 20.4 | 4.4 | 7.9 | 8.0 | 17.6 | 20.0 | 27.2 | 18.2 |
| | BC5-disease | 43.9 | 33.6 | 25.9 | 14.8 | 13.5 | 9.6 | 30.8 | 35.4 | 26.8 | 26.0 |
| | JNLPBA | 35.4 | 40.3 | 42.1 | 17.1 | 19.5 | 8.0 | 47.4 | 51.6 | 17.8 | 31.0 |
| NER | NCBI-disease | 33.5 | 26.8 | 21.8 | 14.5 | 11.7 | 10.4 | 28.0 | 30.8 | 24.6 | 22.5 |
| PICO | EBM-PICO | 3.3 | 39.8 | 45.2 | 1.0 | 9.3 | 2.8 | 37.4 | 41.8 | 18.8 | 22.2 |
| | BioASQ | 21.4 | 18.9 | 15.6 | 33.4 | 10.4 | 16.9 | 16.3 | 17.1 | 7.9 | 17.5 |
| QA | PubMedQA | 19.8 | 53.6 | 57.2 | 28.4 | 1.9 | 20.2 | 31.8 | 48.0 | 4.0 | 29.4 |
| | ChemProt | 36.5 | 58.7 | 57.4 | 11.2 | 36.1 | 6.3 | 57.5 | 61.9 | 12.9 | 37.6 |
| | DDI | 46.7 | 43.7 | 40.4 | 10.0 | 45.1 | 5.7 | 55.3 | 55.8 | 53.8 | 39.6 |
| RE | GAD | 39.0 | 47.6 | 50.9 | 23.8 | 29.5 | 99.8 | 52.5 | 49.5 | 72.5 | 51.7 |
| SS | BIOSSES | 5.5 | 5.9 | 21.0 | 25.4 | 15.4 | 0.0 | 19.1 | 27.0 | 0.0 | 13.2 |
| | **Mean** | 27.9 | 34.2 | 36.4 | 15.2 | 17.4 | 15.6 | 35.7 | 39.5 | 24.0 | 27.3 |

**Supplementary Table S4**. Flex-ECE scores by dataset and model using the hybrid confidence assignment strategy. Only task-model-strategy combinations with a success rate ≥50% are included.

| Task | Dataset | F-XXL | G-3 | G-4 | ML-13 | ML-8 | M-7 | L-8 | Y-34 | Z-7 | Mean |
|---|---|---|---|---|---|---|---|---|---|---|---|
| DC | HoC | 64.6 | 26.4 | 36.1 | 47.5 | 38.6 | 91.8 | 35.4 | 36.5 | 83.5 | 51.2 |
| | BC2GM | 55.0 | 32.9 | 31.1 | 69.1 | 15.4 | 85.4 | 37.0 | 36.3 | 58.1 | 46.7 |
| | BC5-chemical | 40.8 | 21.1 | 20.4 | 63.1 | 8.8 | 89.6 | 16.3 | 19.5 | 60.3 | 37.8 |
| | BC5-disease | 43.9 | 34.9 | 26.0 | 31.3 | 16.6 | 87.0 | 30.3 | 33.8 | 61.3 | 40.6 |
| | JNLPBA | 36.4 | 43.9 | 44.6 | 61.7 | 24.4 | 88.2 | 43.2 | 49.9 | 76.6 | 52.1 |
| NER | NCBI-disease | 50.2 | 27.8 | 22.2 | 32.3 | 12.9 | 87.0 | 32.9 | 29.1 | 62.1 | 39.6 |
| PICO | EBM-PICO | 58.7 | 45.1 | 57.4 | 79.5 | 8.7 | 96.9 | 42.2 | 49.4 | 69.5 | 56.4 |
| | BioASQ | 18.2 | 10.8 | 14.7 | 25.5 | 5.2 | 81.6 | 8.1 | 12.1 | 86.1 | 29.1 |
| QA | PubMedQA | 22.3 | 32.6 | 35.4 | 43.9 | 51.5 | 78.5 | 19.6 | 33.3 | 95.5 | 45.8 |
| | ChemProt | 30.4 | 49.3 | 54.5 | 64.7 | 37.9 | 92.6 | 50.1 | 52.3 | 70.3 | 55.8 |
| | DDI | 47.5 | 42.0 | 40.1 | 62.3 | 43.0 | 93.5 | 47.2 | 49.5 | 31.1 | 50.7 |
| RE | GAD | 20.7 | 45.7 | 50.6 | 23.7 | 25.2 | 0.4 | 43.1 | 41.9 | 10.9 | 29.1 |
| SS | BIOSSES | 74.9 | 11.1 | 8.8 | 17.6 | 14.3 | 97.2 | 15.4 | 18.0 | 100.0 | 39.7 |
| | **Mean** | 43.3 | 32.6 | 34.0 | 47.9 | 23.3 | 82.3 | 32.4 | 35.5 | 66.6 | 44.2 |

#### After Calibration

**Supplementary Table S5.** Flex-ECE scores for each LLM before and after calibration with histogram binning and isotonic regression using a varying amount of calibration examples. Note that if the size of the train split for a particular dataset was smaller than "# Examples", the entire train split was used.

| Task | Dataset | # Examples | Method | F-XXL | G-3 | G-4 | L-8 | M-7 | ML-13 | ML-8 | Y-34 | Z-7 |
|---|---|---|---|---|---|---|---|---|---|---|---|---|
| DC | HoC | 0 | None | 11.3 | 26.4 | 36.1 | 35.4 | 6.1 | 4.6 | 14.5 | 36.5 | 20.1 |
| | | 10 | HB | 21.5 | 25.7 | 24.0 | 18.2 | 0.0 | 13.7 | 27.8 | 41.9 | 0.2 |
| | | | IR | 19.8 | 2.4 | 24.1 | 18.2 | 0.0 | 2.5 | 10.4 | 9.9 | 0.2 |
| | | 50 | HB | 12.5 | 7.3 | 10.3 | 16.1 | 0.0 | 9.7 | 3.1 | 5.2 | 0.2 |
| | | | IR | 9.6 | 1.2 | 7.7 | 8.2 | 0.0 | 7.6 | 2.7 | 7.8 | 0.2 |
| | | 100 | HB | 9.6 | 2.3 | 3.4 | 8.3 | 0.0 | 3.8 | 1.3 | 13.2 | 0.2 |
| | | | IR | 10.0 | 1.8 | 4.4 | 8.3 | 0.0 | 3.7 | 1.6 | 11.2 | 0.2 |
| | | 500 | HB | 5.0 | 2.2 | 2.9 | 6.2 | 0.0 | 1.4 | 0.8 | 3.9 | 0.1 |
| | | | IR | 2.5 | 3.3 | 2.2 | 1.5 | 0.0 | 2.5 | 1.7 | 3.5 | 0.0 |
| | | 1000 | HB | 2.6 | 0.6 | 5.3 | 7.2 | 0.0 | 1.1 | 0.4 | 2.0 | 0.1 |
| | | | IR | 2.2 | 0.1 | 1.0 | 1.1 | 0.0 | 1.2 | 1.5 | 1.9 | 0.0 |
| NER | BC2GM | 0 | None | 28.8 | 32.9 | 31.1 | 37.0 | 9.5 | 8.8 | 11.6 | 36.3 | 25.3 |
| | | 10 | HB | 15.6 | 8.5 | 28.3 | 40.8 | 1.6 | 8.0 | 27.3 | 18.1 | 6.8 |
| | | | IR | 15.4 | 11.4 | 28.3 | 28.0 | 1.6 | 1.3 | 12.9 | 16.5 | 5.1 |
| | | 50 | HB | 9.3 | 9.4 | 12.2 | 13.2 | 0.3 | 0.7 | 9.9 | 9.3 | 10.5 |
| | | | IR | 7.7 | 6.8 | 12.0 | 10.3 | 0.3 | 3.7 | 14.2 | 8.0 | 9.0 |
| | | 100 | HB | 3.8 | 8.5 | 9.7 | 5.4 | 0.1 | 3.6 | 7.2 | 11.8 | 7.5 |
| | | | IR | 5.3 | 6.5 | 10.0 | 4.7 | 0.3 | 4.6 | 12.4 | 9.7 | 7.7 |
| | | 500 | HB | 1.3 | 1.8 | 1.3 | 1.0 | 0.3 | 1.9 | 3.1 | 4.0 | 1.7 |
| | | | IR | 1.5 | 0.7 | 2.1 | 1.0 | 0.2 | 1.7 | 1.6 | 1.6 | 2.4 |
| | | 1000 | HB | 3.0 | 2.3 | 1.0 | 1.3 | 0.4 | 0.0 | 1.8 | 4.1 | 0.9 |
| | | | IR | 1.7 | 0.7 | 1.5 | 0.3 | 0.2 | 0.8 | 0.9 | 1.3 | 1.6 |
| | BC5-chemical | 0 | None | 38.0 | 21.1 | 20.4 | 16.3 | 8.0 | 4.4 | 7.9 | 19.5 | 27.2 |
| | | 10 | HB | 10.5 | 8.9 | 18.6 | 21.9 | 1.1 | 9.9 | 24.0 | 24.0 | 26.2 |
| | | | IR | 14.5 | 11.7 | 16.4 | 8.7 | 0.9 | 4.0 | 16.8 | 23.8 | 26.0 |
| | | 50 | HB | 16.2 | 6.2 | 6.9 | 4.1 | 0.1 | 2.9 | 11.1 | 9.2 | 5.6 |
| | | | IR | 13.5 | 5.9 | 6.3 | 6.8 | 0.1 | 5.5 | 7.8 | 11.5 | 7.7 |
| | | 100 | HB | 9.6 | 4.1 | 4.2 | 3.8 | 0.1 | 1.1 | 8.3 | 11.9 | 3.6 |
| | | | IR | 6.7 | 2.2 | 3.7 | 5.7 | 0.2 | 1.9 | 8.8 | 8.9 | 1.9 |
| | | 500 | HB | 2.9 | 1.9 | 1.1 | 3.6 | 0.3 | 1.9 | 3.3 | 1.8 | 2.2 |
| | | | IR | 1.3 | 2.2 | 2.0 | 3.3 | 0.1 | 1.9 | 4.2 | 1.6 | 1.6 |
| | | 1000 | HB | 2.9 | 1.7 | 0.7 | 0.9 | 0.2 | 1.2 | 0.8 | 1.5 | 2.2 |
| | | | IR | 1.7 | 1.6 | 0.6 | 1.3 | 0.0 | 1.0 | 1.3 | 0.8 | 1.5 |
| | BC5-disease | 0 | None | 43.9 | 34.9 | 26.0 | 30.3 | 9.6 | 14.8 | 13.5 | 33.8 | 26.8 |
| | | 10 | HB | 6.8 | 7.4 | 10.0 | 44.2 | 0.9 | 17.8 | 53.0 | 18.4 | 12.5 |
| | | | IR | 6.7 | 8.6 | 3.1 | 9.5 | 1.9 | 3.2 | 12.0 | 19.4 | 12.1 |
| | | 50 | HB | 3.5 | 6.4 | 1.3 | 10.2 | 0.1 | 13.3 | 5.0 | 8.0 | 5.6 |
| | | | IR | 2.6 | 4.7 | 2.3 | 4.9 | 0.0 | 10.1 | 3.2 | 6.6 | 6.8 |
| | | 100 | HB | 4.3 | 2.7 | 1.8 | 7.6 | 0.3 | 7.9 | 1.7 | 4.2 | 7.8 |
| | | | IR | 4.5 | 4.2 | 1.6 | 4.8 | 0.2 | 9.4 | 3.5 | 4.1 | 8.0 |

| Task | Dataset | # Examples | Method | F-XXL | G-3 | G-4 | L-8 | M-7 | ML-13 | ML-8 | Y-34 | Z-7 |
|---|---|---|---|---|---|---|---|---|---|---|---|---|
| | | 500 | HB | 3.4 | 2.8 | 0.7 | 3.3 | 0.0 | 2.8 | 1.5 | 3.5 | 1.5 |
| | | | IR | 2.6 | 3.1 | 1.5 | 4.5 | 0.0 | 2.5 | 2.4 | 3.4 | 1.1 |
| | | 1000 | HB | 2.8 | 3.4 | 1.5 | 1.8 | 0.1 | 0.7 | 1.1 | 4.2 | 1.2 |
| | | | IR | 2.5 | 0.9 | 2.1 | 1.3 | 0.1 | 1.1 | 4.1 | 3.1 | 1.6 |
| | JNLPBA | 0 | None | 35.4 | 43.9 | 44.6 | 43.2 | 8.0 | 17.1 | 19.5 | 49.9 | 17.8 |
| | | 10 | HB | 40.1 | 15.4 | 14.6 | 19.2 | 0.7 | 5.1 | 32.4 | 12.7 | 10.9 |
| | | | IR | 3.9 | 12.4 | 15.0 | 5.7 | 0.3 | 5.1 | 26.8 | 8.9 | 2.1 |
| | | 50 | HB | 6.0 | 1.9 | 3.8 | 14.9 | 0.2 | 0.7 | 10.4 | 6.8 | 2.5 |
| | | | IR | 6.5 | 0.3 | 4.3 | 13.6 | 0.5 | 1.1 | 11.4 | 3.1 | 4.2 |
| | | 100 | HB | 2.7 | 2.9 | 0.4 | 9.2 | 0.2 | 5.2 | 9.4 | 2.3 | 0.4 |
| | | | IR | 5.7 | 2.1 | 1.1 | 9.2 | 0.1 | 1.1 | 7.5 | 1.3 | 4.2 |
| | | 500 | HB | 2.1 | 1.2 | 1.6 | 6.4 | 0.5 | 2.0 | 5.0 | 2.1 | 1.1 |
| | | | IR | 1.0 | 0.9 | 1.3 | 4.8 | 0.7 | 2.1 | 3.7 | 2.1 | 0.8 |
| | | 1000 | HB | 5.3 | 1.3 | 1.3 | 4.3 | 0.5 | 1.9 | 2.5 | 1.7 | 1.1 |
| | | | IR | 0.8 | 2.5 | 1.7 | 3.5 | 0.7 | 1.5 | 3.0 | 1.9 | 1.3 |
| | NCBI-disease | 0 | None | 33.5 | 27.8 | 22.2 | 32.9 | 10.4 | 14.5 | 11.7 | 29.1 | 24.6 |
| | | 10 | HB | 20.4 | 21.8 | 0.3 | 25.6 | 0.7 | 39.9 | 48.2 | 32.0 | 15.2 |
| | | | IR | 20.3 | 22.0 | 0.3 | 22.9 | 0.7 | 14.3 | 1.4 | 29.4 | 16.1 |
| | | 50 | HB | 11.2 | 7.2 | 0.7 | 15.3 | 2.2 | 10.2 | 10.2 | 3.8 | 2.9 |
| | | | IR | 9.1 | 6.4 | 2.5 | 12.3 | 2.5 | 10.7 | 9.1 | 4.4 | 3.2 |
| | | 100 | HB | 8.3 | 2.2 | 2.5 | 8.8 | 1.3 | 1.4 | 2.3 | 7.3 | 2.2 |
| | | | IR | 6.7 | 1.6 | 3.1 | 6.6 | 1.4 | 3.2 | 3.6 | 4.4 | 1.4 |
| | | 500 | HB | 3.8 | 2.1 | 1.1 | 5.3 | 0.2 | 5.1 | 9.2 | 1.7 | 4.8 |
| | | | IR | 3.9 | 4.4 | 3.7 | 6.8 | 0.1 | 4.2 | 7.7 | 3.2 | 3.8 |
| | | 1000 | HB | 5.5 | 4.6 | 1.3 | 6.5 | 0.0 | 2.7 | 4.5 | 1.9 | 4.2 |
| | | | IR | 4.4 | 4.7 | 2.8 | 4.4 | 0.1 | 2.9 | 3.1 | 1.1 | 2.3 |
| PICO | EBM-PICO | 0 | None | 3.3 | 45.1 | 57.4 | 42.2 | 2.8 | 1.0 | 9.3 | 49.4 | 18.8 |
| | | 10 | HB | 2.0 | 8.3 | 12.6 | 10.2 | 0.1 | 0.2 | 0.5 | 10.6 | 1.6 |
| | | | IR | 1.8 | 8.1 | 12.6 | 10.9 | 0.1 | 0.2 | 1.8 | 3.4 | 1.6 |
| | | 50 | HB | 0.8 | 4.5 | 3.9 | 3.4 | 0.1 | 1.2 | 1.0 | 3.6 | 1.3 |
| | | | IR | 0.6 | 3.1 | 3.3 | 4.6 | 0.0 | 1.1 | 1.1 | 3.0 | 0.4 |
| | | 100 | HB | 0.7 | 4.2 | 4.7 | 2.0 | 0.1 | 0.9 | 1.1 | 3.2 | 0.3 |
| | | | IR | 0.4 | 3.6 | 4.0 | 1.9 | 0.1 | 0.7 | 0.8 | 2.9 | 0.7 |
| | | 500 | HB | 0.9 | 4.1 | 5.3 | 4.2 | 0.4 | 0.7 | 0.7 | 2.5 | 1.2 |
| | | | IR | 0.7 | 4.0 | 4.7 | 3.7 | 0.4 | 0.6 | 0.4 | 2.4 | 1.4 |
| | | 1000 | HB | 0.7 | 4.6 | 3.9 | 4.4 | 0.3 | 0.5 | 1.1 | 2.1 | 1.2 |
| | | | IR | 0.6 | 4.2 | 3.7 | 3.8 | 0.3 | 0.4 | 0.8 | 2.1 | 1.4 |
| QA | BioASQ | 0 | None | 21.4 | 10.8 | 14.7 | 8.1 | 16.9 | 33.4 | 10.4 | 12.1 | 7.9 |
| | | 10 | HB | 25.5 | 25.7 | 16.4 | 20.5 | 0.0 | 20.0 | 20.7 | 37.1 | 5.0 |
| | | | IR | 13.5 | 11.7 | 16.4 | 10.0 | 0.0 | 23.2 | 12.9 | 33.7 | 2.9 |
| | | 50 | HB | 14.1 | 13.1 | 9.3 | 16.7 | 0.0 | 16.5 | 8.1 | 10.9 | 3.6 |
| | | | IR | 15.0 | 11.9 | 8.7 | 14.6 | 0.0 | 21.3 | 7.2 | 3.8 | 4.7 |
| | | 100 | HB | 8.1 | 7.5 | 7.5 | 10.0 | 0.0 | 19.7 | 5.9 | 2.0 | 0.5 |
| | | | IR | 5.3 | 6.5 | 7.5 | 9.2 | 0.0 | 18.4 | 5.7 | 3.2 | 3.8 |
| | | 500 | HB | 6.8 | 5.2 | 4.8 | 5.3 | 0.0 | 14.7 | 6.1 | 4.8 | 1.2 |

| Task | Dataset | # Examples | Method | F-XXL | G-3 | G-4 | L-8 | M-7 | ML-13 | ML-8 | Y-34 | Z-7 |
|---|---|---|---|---|---|---|---|---|---|---|---|---|
| | | | IR | 7.0 | 5.2 | 8.5 | 5.8 | 0.0 | 14.5 | 4.9 | 5.9 | 2.6 |
| | | 1000 | HB | 7.1 | 4.3 | 4.8 | 4.8 | 0.0 | 15.1 | 3.2 | 2.1 | 1.4 |
| | | | IR | 6.3 | 4.4 | 8.0 | 5.5 | 0.0 | 15.2 | 4.2 | 4.2 | 2.7 |
| | PubMedQA | 0 | None | 19.8 | 32.6 | 35.4 | 19.6 | 20.2 | 28.4 | 1.9 | 33.3 | 4.0 |
| | | 10 | HB | 45.2 | 29.8 | 27.2 | 39.0 | 0.0 | 29.8 | 4.4 | 40.2 | 0.2 |
| | | | IR | 38.9 | 17.8 | 17.4 | 16.6 | 0.0 | 22.1 | 4.4 | 27.4 | 0.2 |
| | | 50 | HB | 8.6 | 11.6 | 13.4 | 9.1 | 0.0 | 5.4 | 4.0 | 9.8 | 0.2 |
| | | | IR | 5.2 | 5.6 | 6.1 | 4.7 | 0.0 | 7.3 | 3.4 | 8.3 | 0.2 |
| | | 100 | HB | 6.8 | 6.0 | 6.4 | 5.6 | 0.0 | 6.1 | 3.2 | 7.6 | 0.2 |
| | | | IR | 5.8 | 1.2 | 8.0 | 2.8 | 0.0 | 7.9 | 3.0 | 7.8 | 0.2 |
| | | 500 | HB | 7.0 | 3.8 | 2.6 | 1.0 | 0.0 | 2.5 | 2.2 | 3.4 | 0.2 |
| | | | IR | 5.4 | 1.8 | 1.9 | 2.7 | 0.0 | 2.7 | 1.6 | 4.7 | 0.2 |
| | | 1000 | HB | 7.0 | 3.8 | 2.6 | 1.0 | 0.0 | 2.5 | 2.2 | 3.4 | 0.2 |
| | | | IR | 5.4 | 1.8 | 1.9 | 2.7 | 0.0 | 2.7 | 1.6 | 4.7 | 0.2 |
| RE | ChemProt | 0 | None | 36.5 | 49.3 | 54.5 | 50.1 | 6.3 | 11.2 | 36.1 | 52.3 | 12.9 |
| | | 10 | HB | 29.6 | 18.6 | 9.2 | 13.4 | 0.0 | 3.2 | 13.9 | 30.5 | 17.9 |
| | | | IR | 26.8 | 2.5 | 9.5 | 5.6 | 0.0 | 3.2 | 7.8 | 13.6 | 17.9 |
| | | 50 | HB | 13.5 | 6.1 | 11.5 | 3.8 | 0.0 | 0.2 | 14.9 | 20.0 | 10.2 |
| | | | IR | 4.1 | 7.5 | 15.1 | 5.7 | 0.0 | 1.7 | 14.3 | 13.7 | 11.8 |
| | | 100 | HB | 4.5 | 3.5 | 13.9 | 2.3 | 0.0 | 6.7 | 11.4 | 3.8 | 2.8 |
| | | | IR | 2.7 | 4.8 | 13.9 | 3.9 | 0.0 | 5.3 | 12.7 | 4.6 | 3.4 |
| | | 500 | HB | 1.6 | 3.4 | 4.6 | 4.2 | 0.0 | 1.2 | 1.9 | 2.1 | 3.3 |
| | | | IR | 2.6 | 1.2 | 5.3 | 3.1 | 0.0 | 1.6 | 2.9 | 4.3 | 1.0 |
| | | 1000 | HB | 2.3 | 0.4 | 3.8 | 2.2 | 0.0 | 0.5 | 0.5 | 1.2 | 1.0 |
| | | | IR | 1.9 | 1.3 | 4.7 | 2.3 | 0.0 | 0.8 | 2.1 | 2.7 | 2.4 |
| | DDI | 0 | None | 46.7 | 42.0 | 40.1 | 47.2 | 5.7 | 10.0 | 45.1 | 49.5 | 53.8 |
| | | 10 | HB | 26.5 | 36.0 | 27.1 | 39.5 | 0.0 | 54.4 | 51.1 | 28.0 | 12.3 |
| | | | IR | 24.8 | 38.2 | 27.9 | 26.6 | 0.0 | 18.4 | 17.3 | 8.2 | 0.4 |
| | | 50 | HB | 3.6 | 4.6 | 3.3 | 14.7 | 0.0 | 9.5 | 4.7 | 7.2 | 8.9 |
| | | | IR | 2.9 | 6.1 | 3.6 | 3.8 | 0.0 | 8.1 | 5.7 | 4.3 | 6.3 |
| | | 100 | HB | 6.0 | 0.8 | 7.3 | 4.3 | 0.0 | 4.5 | 4.5 | 7.7 | 4.9 |
| | | | IR | 4.3 | 1.9 | 5.1 | 4.2 | 0.0 | 3.6 | 5.4 | 8.8 | 4.0 |
| | | 500 | HB | 1.1 | 4.2 | 4.7 | 2.9 | 0.0 | 5.4 | 3.1 | 4.8 | 3.5 |
| | | | IR | 0.9 | 4.0 | 4.7 | 2.7 | 0.0 | 4.7 | 2.4 | 4.4 | 2.9 |
| | | 1000 | HB | 1.1 | 2.4 | 4.7 | 4.7 | 0.0 | 3.5 | 5.4 | 2.9 | 4.1 |
| | | | IR | 1.1 | 2.3 | 2.9 | 3.8 | 0.0 | 4.1 | 5.1 | 3.0 | 3.9 |
| | GAD | 0 | None | 39.0 | 45.7 | 50.6 | 43.1 | 99.8 | 23.8 | 29.5 | 41.9 | 72.5 |
| | | 10 | HB | 27.1 | 30.2 | 21.7 | 26.6 | 0.0 | 52.8 | 34.4 | 50.6 | 17.8 |
| | | | IR | 30.4 | 30.0 | 21.7 | 19.2 | 0.0 | 8.4 | 39.1 | 40.8 | 12.2 |
| | | 50 | HB | 24.1 | 26.7 | 21.7 | 24.0 | 0.0 | 13.9 | 22.8 | 24.0 | 17.8 |
| | | | IR | 24.6 | 26.7 | 21.7 | 27.0 | 0.0 | 8.4 | 22.5 | 22.2 | 8.2 |
| | | 100 | HB | 13.8 | 17.1 | 14.7 | 19.0 | 0.0 | 15.1 | 15.2 | 13.9 | 5.9 |
| | | | IR | 13.4 | 15.3 | 15.2 | 18.8 | 0.0 | 7.4 | 14.2 | 16.6 | 4.2 |
| | | 500 | HB | 0.2 | 1.2 | 0.1 | 3.8 | 0.0 | 2.0 | 2.3 | 0.8 | 3.1 |
| | | | IR | 0.0 | 1.3 | 0.9 | 8.6 | 0.0 | 0.6 | 1.9 | 0.3 | 1.6 |

| Task | Dataset | # Examples | Method | F-XXL | G-3 | G-4 | L-8 | M-7 | ML-13 | ML-8 | Y-34 | Z-7 |
|---|---|---|---|---|---|---|---|---|---|---|---|---|
| | | 1000 | HB | 0.0 | 0.5 | 1.2 | 6.9 | 0.0 | 4.9 | 1.7 | 1.8 | 3.3 |
| | | | IR | 0.2 | 0.4 | 1.2 | 7.2 | 0.0 | 1.8 | 1.7 | 1.3 | 1.7 |
| SS | BIOSSES | 0 | None | 5.5 | 11.1 | 8.8 | 15.4 | 0.0 | 25.4 | 15.4 | 18.0 | 0.0 |
| | | 10 | HB | 5.2 | 27.6 | 11.8 | 13.4 | 0.0 | 25.4 | 38.7 | 26.9 | 0.0 |
| | | | IR | 4.9 | 25.9 | 11.7 | 13.0 | 0.0 | 27.4 | 32.7 | 25.0 | 0.0 |
| | | 50 | HB | 5.2 | 5.8 | 13.2 | 10.5 | 0.0 | 9.7 | 4.4 | 5.6 | 0.0 |
| | | | IR | 0.9 | 5.9 | 4.9 | 14.1 | 0.0 | 10.1 | 6.2 | 7.1 | 0.0 |
| | | 100 | HB | 2.1 | 2.3 | 6.8 | 12.0 | 0.0 | 14.1 | 6.2 | 4.4 | 0.0 |
| | | | IR | 0.8 | 10.0 | 4.6 | 11.2 | 0.0 | 11.8 | 4.0 | 7.9 | 0.0 |
| | | 500 | HB | 2.1 | 2.3 | 6.8 | 12.0 | 0.0 | 14.1 | 6.2 | 4.4 | 0.0 |
| | | | IR | 0.8 | 10.0 | 4.6 | 11.2 | 0.0 | 11.8 | 4.0 | 7.9 | 0.0 |
| | | 1000 | HB | 2.1 | 2.3 | 6.8 | 12.0 | 0.0 | 14.1 | 6.2 | 4.4 | 0.0 |
| | | | IR | 0.8 | 10.0 | 4.6 | 11.2 | 0.0 | 11.8 | 4.0 | 7.9 | 0.0 |
